# Cohesin extrudes chromatin loop unidirectionally through two modes of mechanisms in human cells

**DOI:** 10.64898/2026.05.01.722300

**Authors:** Ping Wang, Lingyun Meng, Linghan Jiang, Yang Li, Jiaxiang Huang, Tiancheng Yu, Haoxi Chai, Minji Kim, Xiaotao Wang, Yijun Ruan

## Abstract

Cohesin is crucial for establishing 3D genome organization and functions. However, the *in vivo* mechanism of cohesin-mediated extrusion remains unsettled. Here, we investigated extrusion directionality using integrated genome mapping coupled with perturbation in human cells. By defining cohesin loading at NIPBL binding loci and anchoring at CTCF sites, we systematically characterized the aggregated and single-molecule trajectories of extrusion “stripes” and contact “dots” from and between individual cohesin loading and anchoring points. Our analyses delineated two dynamic modes of *in vivo* mechanism in which NIPBL/cohesin initiates extrusion at its loading sites in a ‘one-sided’ manner; after encountering CTCF, CTCF/cohesin switches its extrusion direction backward and robustly extrude DNA unidirectionally until completing the convergent CTCF loop. Surprisingly, depleting CTCF resulted substantial increase of NIPBL/cohesin-mediated extrusion activity, which is further validated by *in silico* perturbation. These findings reveal novel mechanistic insights into how cohesin and CTCF orchestrate cohesin-mediated loop extrusion in vivo.

## INTRODUCTION

The three-dimensional (3D) genome organization in human cells is characterized by chromatin interaction loops as its basic topological units,^1–4^ which further aggregate into higher-order topologically associating domains (TADs),^5–7^ compartments,^8–11^ and chromosome territories within the nuclear space.^12–14^ This multi-level 3D genome organization provides a topological framework to modulate temporal nuclear functions including gene transcription ^15–18^ and DNA replication.^19–21^ In particular, the ring-shaped cohesin protein complex SMC (Structural Maintenance of Chromosomes) proteins play key roles through a process called loop extrusion to establish the topological structures of the genome.^22–30^ Several chromatin architecture proteins are also involved in regulating the process of chromatin loop formation, including NIPBL and WAPL for cohesin loading and releasing, respectively, and CTCF for defining the boundaries of cohesin-mediated extrusion. Functional perturbation experiments rapidly depleting cohesion,^31–35^ CTCF, ^26,36–39^ NIPBL ^40,41^ and WAPL ^33,42–45^ have validated their respective roles and provided a broader view of cohesin-mediated chromatin extrusion during the interphase of cells. However, the detailed mechanism by which cohesin extrudes the DNA remains unsettled as summarized below.

Most of our mechanistic understandings of cohesin-mediated loop extrusion are derived from *in vitro* biochemical reconstitution and protein structural conformation studies.^28,46–49^ It has been suggested that cohesin is loaded onto the DNA strand at the NIPBL binding sites in transcriptionally active regions,^50–53^ and the NIPBL/cohesin complex extrudes DNA symmetrically in a “two-sided” bidirectional manner ^28^ or through a “swing and clamp” mechanism.^47,54^ However, several recent *in vitro* studies using single-molecule experiments suggested that the human cohesin does not reel in DNA from both sides as previously reported; instead, it extrudes the DNA asymmetrically.^48,55^ Thus, whether cohesin extrudes DNA symmetrically or asymmetrically is still controversial in the field even *in vitro*.

More importantly, the questions regarding how cohesin extrudes DNA *in vivo* are still wide open. Although it has been generally perceived that cohesin could extrude DNA two-sided symmetrically from its loading sites until anchored at CTCF sites,^22,24,28,46,56,57^ where the stripe signals are profoundly recognized in Hi-C data,^58^ the actual directionality of cohesin extrusion *in vivo* remains debatable. Theoretically, if the two-sided symmetric extrusion model is applicable *in vivo*, the 2D contact profiles from Hi-C mapping data should be readily displaying a strong signal perpendicular to the diagonal line of the square heat map. Such pattern has only been explicitly observed in bacteria ^25,59^ and have rarely been seen in mammalian cells.^60^ However, several recent studies have reported such perpendicular mapping signals, referred to as chromatin ‘jets’ ^61^ or ‘fountain’,^62^ in Hi-C data derived from either quiescent or manipulated mouse cells. Concerns are that the 37 handpicked ‘jets’ may exist only in quiescent cells as they have not been seen in other mouse cells, and that the observed ‘fountains’ are likely artifacts due to depleted functions in the perturbed mouse cells.^60^ Although similar chromatin fountain structures had also been reported in zebrafish and *C. elegans*, ^63,64^ a common characteristic of the reported jets or fountains in various species is their unusual large size in the range of megabases, which is clearly out of the range of cohesin-mediated chromatin loops with convergent CTCF anchors (averaging 200 kb),^65^ indicating that the observed fountains may arise from other nuclear activities such as DNA replication.^66^ Even though the chromatin architectural ‘stripes’ rooted from CTCF binding sites had been observed at thousands of loci across many mammalian cell lines,^58,67^ the model of cohesin-mediated chromatin looping *in vivo* remains largely unclear,^60^ likely due to the lack of mapping resolution and specificity for the identification of cohesin-mediated extrusion events in human cells.

To address the *in vivo* mechanism of cohesin-mediated chromatin loop extrusion, we applied an integrative genome-wide mapping strategy that included ChIA-Drop and ChIA-PET with enriched specificity for cohesin- and CTCF-associated chromatin interactions and Micro-C for deep coverage and high-resolution mapping. Focusing on a large set of well-defined cohesin loading and anchoring loci in the context of convergent CTCF loops, we comprehensively characterized cohesin-mediated extrusion trajectories with single-molecule resolution and targeted enrichment for cohesin and CTCF specificity along with perturbation experiments for functional interpretation. Our results provide convincing evidence to suggest that cohesin extrudes DNA unidirectionally from its initial loading sites with NIPBL and at its anchoring sites with CTCF until completing a chromatin looping process with convergent CTCF anchors.

## RESULTS

### ChIA-Drop single-molecule mapping of multiplex interactions with protein specificity

To unravel the intricate details of the dynamic process of chromatin looping trajectory *in vivo*, we analyzed the human genome by performing ChIA-Drop,^68^ a ligation-free method for mapping multiplex chromatin interactions with single-molecule resolution (**Figure 1A**). ChIA-Drop data from B-lymphoblastoid cells (GM19239, GM12878) were processed using the ChIA-DropBox pipeline ^69,70^ and MIA-Sig algorithm^68,71^ to identify chromatin looping complexes containing multiple chromatin fragments indexed with the same droplet barcode (see **Methods**). The processed ChIA-Drop data were visualized in 2D contact profiles for pairwise interactions and displayed in browser views for multiplex chromatin looping complexes that are largely comparable to those of the Hi-C^5^ and SPRITE,^72^ confirming that the ChIA-Drop data derived from human cells are of high quality (**Figure 1B-C, Figure S1A-G**).

**Figure 1.**
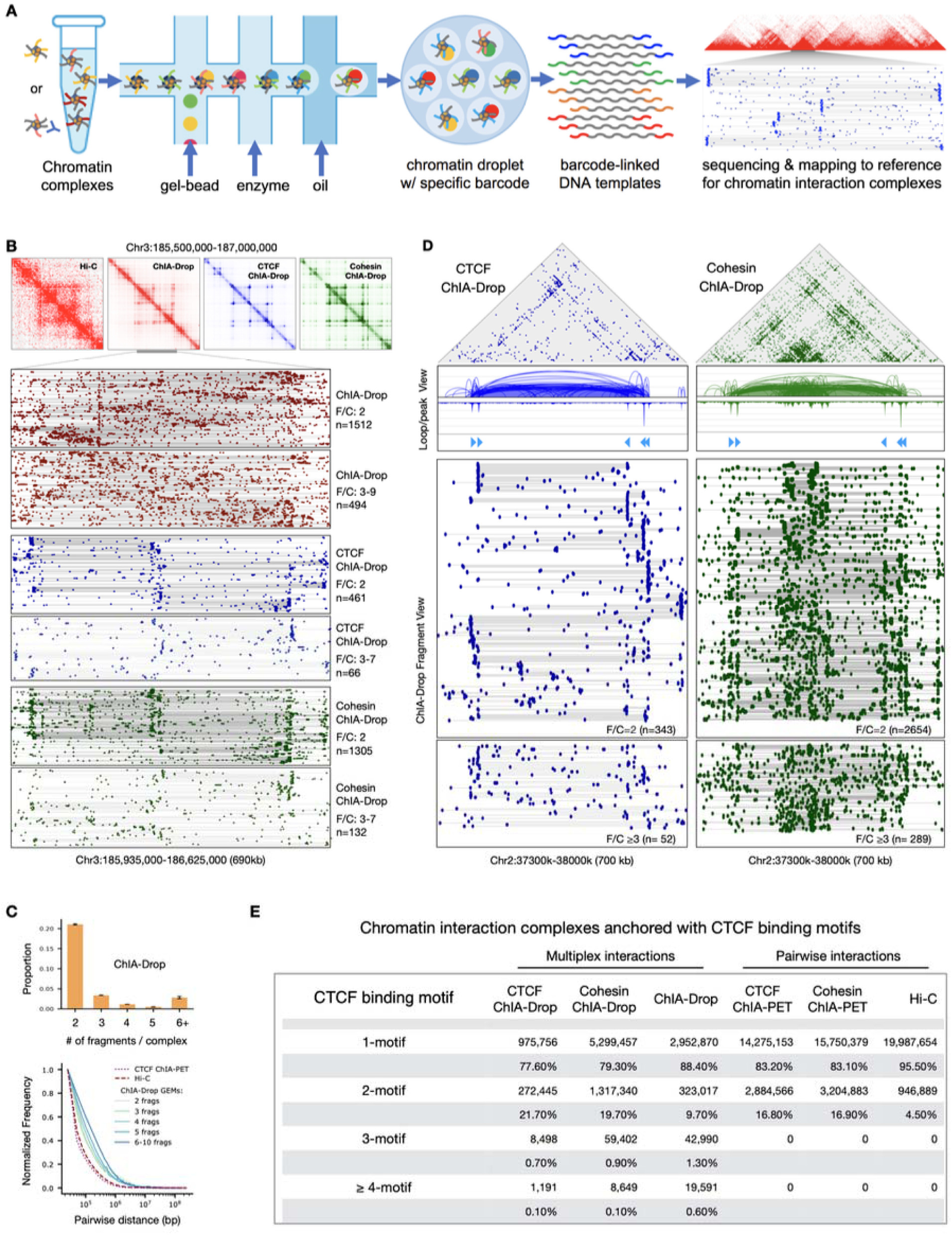
ChIA-Drop data mapping multiplex chromatin interactions mediated by CTCF and cohesion. (**A**) A schematic of ChIA-Drop, which encapsulates ChIP-enriched or non-enriched samples of chromatin interaction complexes in individual droplets with unique barcodes for obtaining single-molecule chromatin interactions via DNA sequencing and mapping analysis. (**B**) Example views of 2D contact maps in comparison of Hi-C and ChIA-Drop data (top panel). The fragment views of multiplex chromatin interactions in ChIA-Drop data with 2 fragments per complex (F/C) and 3-9 F/C are presented (bottom panel). The numbers (n) of complexes in each category are provided. (**C**) Bar chart of ChIA-Drop data in multiplexity (top panel) and curves of contact frequency for pairwise distances of ChIA-Drop chromatin complexes decomposed by the number of fragments per complex (green to blue lines) in comparison with Hi-C and CTCF ChIA-PET data in red dashed and dotted lines, respectively (bottom panel). (**D**) Examples of CTCF- and cohesin-enriched ChIA-Drop data in 2D contacts (top panel), loop/peak (middle panel), and fragment views (bottom panel). CTCF binding motifs and fragment counts per chromatin complex (F/C) are provided. (**E**) Table of chromatin complexes associated with the number of CTCF binding motifs in various datasets.

It is known that most *bona fide* chromatin interactions are mediated by protein factors known for either structural maintenance or functional capacity. Specifically, CTCF and cohesin are known involved in the formation of chromatin loops. To investigate the interplay between CTCF and cohesin in shaping chromatin topology with single-molecule mapping resolution, we introduced a chromatin immunoprecipitation (ChIP) step to the ChIA-Drop protocol and probed CTCF and cohesin complex by targeting its subunits SMC1A and RAD21, respectively, to generate ChIA-Drop data in GM12878 cells (**Table S1**). The high correlation between replicates of the ChIP-enriched ChIA-Drop data by CTCF, SMC1A and RAD21 indicates that the data are reproducible and are of high fidelity (**Figure S1H**). Considering that both SMC1A and RAD21 are subunits of cohesin and given the high correlation between SMC1A and RAD21 ChIA-Drop data (**Figure S1H**), we merged these datasets to constitute the cohesin ChIA-Drop data. All ChIA-Drop data exhibited higher-order TAD structures and boundary insulation capacity (**Figure S1I**) comparable to previously reported Hi-C data.^5^ In addition, the CTCF and cohesin enriched datasets (ChIA-Drop and ChIA-PET) displayed much clearer edges (stripes) of TAD structures than the non-enriched data (ChIA-Drop and Hi-C) in 2D contact maps (**Figure S1I**), demonstrating the specificity of chromatin features mediated by the CTCF and cohesin. The browser views of CTCF and cohesin ChIA-Drop data showed additional detailed presentation of enriched chromatin fragments mapped at the CTCF binding motifs and the anchors of chromatin loops with single molecule resolution (**Figure 1D**). The numbers of chromatin fragments included in a chromatin looping complex overlapping with CTCF sites provide characteristic features of the ChIA-Drop and related datasets (**Figure 1E**). Overall, most (around 80%) of the chromatin contacts in all datasets contained only one fragment that was aligned to a CTCF site, indicating the capturing of potential ongoing (incomplete) looping events. Specifically, in the CTCF and cohesin ChIA-Drop data, a significant portion (around 20%) of the chromatin looping complexes possessed fragments mapped to two CTCF sites, likely representing the completed loop structure anchored at two CTCF sites. A small fraction (about 1%), but still large numbers (tens of thousands), contained 3 or more CTCF sites, suggesting the presence of multiplexed clusters of CTCF loops (**Figure 1E**).

It is noteworthy that besides the CTCF and cohesin ChIA-Drop mapping data highly aligned at CTCF sites, abundant cohesin ChIA-Drop mapping signals were found distal from CTCF sites within CTCF/cohesin looping domain (**Figure 1D**, right panel), reflecting potential CTCF-independent and cohesin-specific chromatin looping. This observation prompted us to address an unresolved question regarding the *in vivo* mechanism of cohesin-mediated chromatin loop extrusion and the interplays between cohesin and CTCF in human cells.

### Most cohesin loading sites are encapsulated in chromatin looping domains and away from CTCF binding sites

To comprehensively investigate the models of cohesin-mediated chromatin extrusion *in vivo*, we sought first to systematically identify cohesin loading sites in GM12878 cells. While cohesin is anchored at CTCF binding sites of chromatin loops, where exactly cohesin loads onto the DNA to start loop extrusion is less clear. Although NIPBL is known as a cohesin loading factor, the actual NIPBL binding sites have not been fully characterized^73^ as there is no known NIPBL binding motif reported thus far. Therefore, we generated high-quality NIPBL-seq data and analyzed additional ChIP-seq datasets of several protein factors known to facilitate cohesin’s function in chromatin extrusion and histone markers associated with active transcription (**STAR Methods**). While the majority of the cohesin sites is co-localized with CTCF as potential cohesin anchoring sites, more than one third of the cohesin sites are devoid of CTCF and are highly associated with NIPBL, thereby constituting potential “cohesin loading” sites (**Figure 2A, S2A**). It is noteworthy that there is a small fraction of the NIPBL sites that also co-localized with CTCF/cohesin loci, indicating that a subset of cohesin could also directly load nearby CTCF sites. In addition, all cohesin loading sites are highly associated with active transcriptional signals (RNAPII, H3K27ac and H3K4me1), whereas most of the CTCF/cohesin anchoring sites are largely correlated with insulating and repressive features (**Figure 2A, S2B**). Intriguingly, WAPL, known as a cohesin releasing factor and in competing with NIPBL for cohesin, is co-localized with cohesin at both loading and anchoring loci, indicating that WAPL is somehow constantly associated with cohesin to balance its extrusion function.^43^

**Figure 2.**
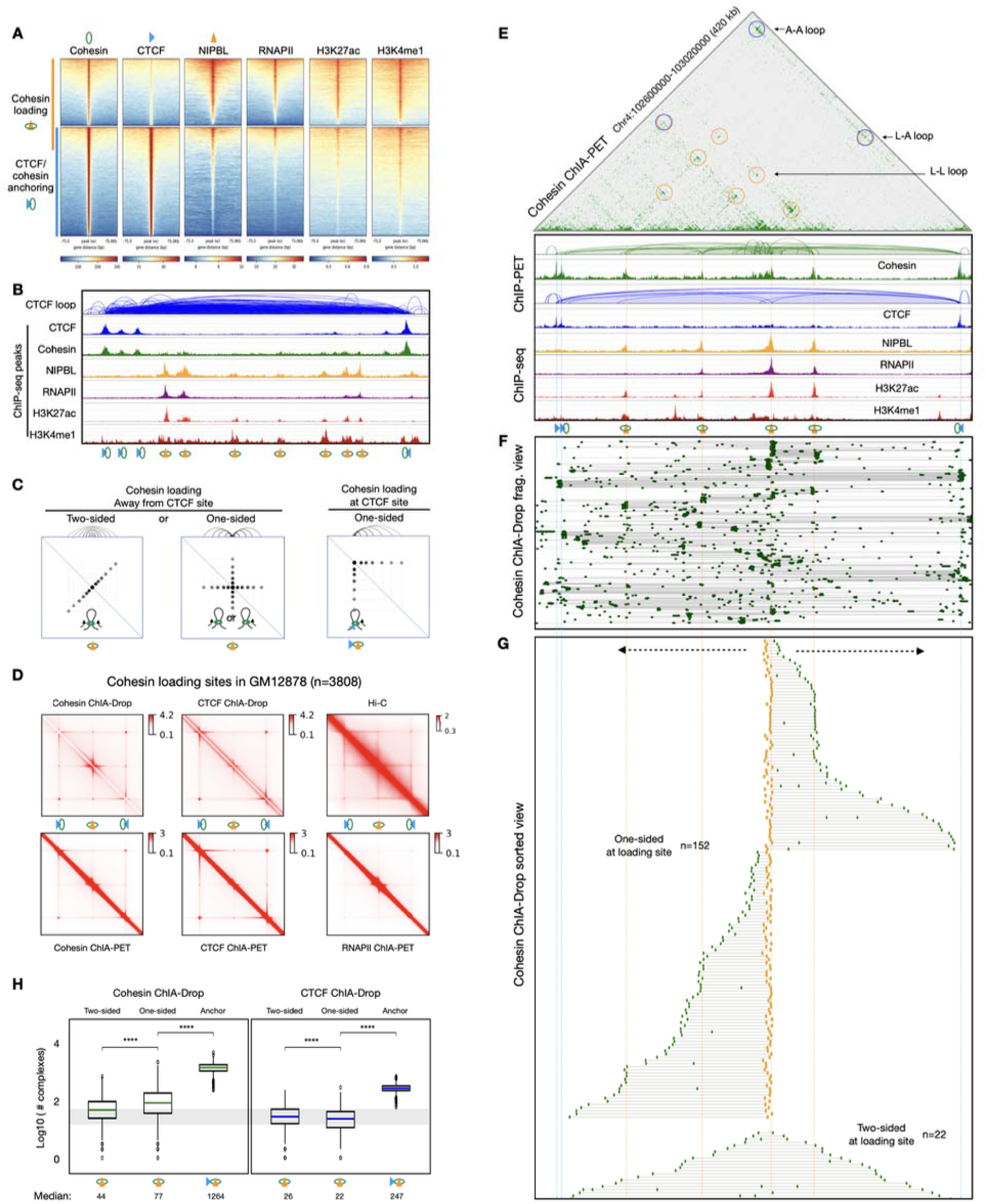
One-sided loop extrusion from cohesin loading sites. (**A**) Aggregated heatmaps of cohesin and CTCF binding peak intensity at their binding sites derived from ChIP-seq datasets. The corresponding binding peak signals of NIPBL, RNAPII, H3K27ac, and H3K4me1 ChIP-seq data are also included. Cohesin loading with NIPBL sites and cohesin anchoring with CTCF sites are demarcated. Most of cohesin loading sites are not overlapped with CTCF binding sites. (**B**) An example locus with a convergent CTCF loop encompassing cohesin loading and anchoring sites as characterized by ChIA-PET and ChIP-seq data. (**C**) Diagrams of three theoretical scenarios of cohesin-mediated chromatin extrusion at cohesin loading (NIPBL binding) sites: two-sided symmetric extrusion would display a stripe perpendicular to the diagonal line (left panel); one-sided extrusion would reel in DNA in either direction and result a cross-shaped stripe pattern in the center (middle panel); if a cohesin loads nearby a CTCF binding site, it would show a stripe that is 45 degrees to the diagonal line (right panel). (**D**) The 2D contact matrices from aggregated chromatin interactions centered at cohesin loading sites (n=3,808) localized within canonical convergent CTCF/cohesin loops in GM12878 cells. Left panel: cohesin enriched ChIA-Drop (top) and ChIA-PET (bottom) data. Middle panel: CTCF enriched ChIA-Drop (top) and ChIA-PET (bottom) data. Right panel: Hi-C (top) and RNAPII enriched ChIA-PET (bottom) data. (**E**) An example locus. Top panel: cohesin ChIA-PET 2D contact map with cohesin loading sites derived cohesin-mediated stripes and contact dots demarcating potential loading-to-loading (L-L), loading-to-anchor (L-A), and anchor-to-anchor (A-A) loops. Bottom panel: browser views of ChIA-PET and ChIP-seq data identified chromatin loops and cohesin loading and anchoring sites. (**F**) Fragment view of multiplex chromatin interactions in 32ohesion ChIA-Drop data in the same region as in **E**. (**G**) Sorted fragment view for the same region as in **F** showing single-molecule mapping trajectories of cohesin-mediated extrusion from one of the loading sites in this looping domain in possible one-sided and two-sided manners. The DNA fragments aligned with the loading site are highlighted in yellow. Extrusion directions are indicated by dotted arrows. The numbers (n) of single-molecule mapping events are provided. (**H**) Boxplots of the numbers of ChIA-Drop mapped chromatin complexes observed at cohesin loading sites in cohesin ChIA-Drop (left panel) and CTCF ChIA-Drop (right panel) data. Note: **** *p*-values (< 2.22e-16) were computed between pairs of datasets using the two-sided Mann-Whitney U test.

We then focused on the cohesin loading sites in the context of canonical CTCF chromatin loops characteristically with a pair of convergent CTCF binding motifs at loop anchoring points. A well-defined cohesin loading locus is characterized by having distinctive NIPBL binding peak with recognizable cohesin occupancy signals and supported by RNAPII, H3K27ac and H3K4me1. Together, by incorporating multiple lines of evidence, we identified 4878 cohesin-loading sites that are encapsulated by CTCF/cohesin-anchored loops, of which 3808 sites are located inside the loops and distal to loop anchoring sites, while 1070 sites are proximal to CTCF sites in GM12878 cells (see **Methods**). Note that a canonical CTCF/cohesin-anchored chromatin looping domain could have one or multiple cohesin-loading sites (**Figure 2B, S2C**).

### Cohesin extrude chromatin loops from its loading-sites *in vivo* in a one-sided model

It is generally viewed that cohesin extrudes DNA in a two-sided symmetrical manner mostly suggested by *in vitro* biophysical experiments with single molecule data^28^ and computational simulation,^57^ even though this view has been challenged by recent *in vitro* reports.^55^ However, how cohesin extrudes DNA *in vivo* from Hi-C mapping data remains vague. With the well-defined cohesin loading sites, it is now possible to re-analyze the publicly available mapping data and our own ChIA-Drop data to study *in vivo* extrusion with single-molecule resolution.

In 2D contact matrix, a mapping feature of two-sided extrusion would theoretically create a contact signal perpendicular to the diagonal line from a cohesin-loading site in a 2D contact square heatmap with the cohesin loading-sites in the center (**Figure 2C**, left panel), similar to what had been observed of DNA folding configuration in bacteria.^25,59^ However, this two-sided mapping pattern had not been seen in most available mammalian 3D genome mapping data,^60^ except for the chromatin “jet” reported in small numbers (n=37) of cohesin-associated loci in quiescent or perturbed mouse cells.^61,74^ An alternative scenario would involve cohesin translocating in a one-sided manner and moving toward left or right without directional preference from a given starting point, thereby forming a ‘cross-shaped’ pattern of ensemble mapping signals in a 2D contact heatmap (**Figure 2C**, middle panel). On the contrary, at a CTCF binding site, cohesin is known to extrude DNA unidirectionally in concordance with the orientation of CTCF binding motif, which were often observed in 2D contact maps as a ‘stripe’ signal pattern (**Figure 2C**, right panel) in mammalian cells.^58,67^ Nevertheless, how exactly the cohesin-mediated actions from its loading to its anchoring sites is still largely unsettled.

To examine the directionality of cohesin-mediated extrusion *in vivo*, we focused on the 3808 well-defined cohesin loading sites that are encapsulated in canonical chromatin loops and are distal from the loop anchoring sites with convergent CTCF binding motifs (see **Methods**). As shown in the results of aggregation peak analysis, the square plot of the aggregated cohesin ChIA-Drop data exhibited a distinct ‘cross-shaped’ signal pattern at the cohesin-loading site along with the typical ‘stripes’ rooted from the paired convergent anchoring sites (**Figure 2D**, top left panel), which clearly lacked any observable ‘jet’ signals ^61^ to suggest a possible two-sided extrusion pattern. In addition, besides the strong looping “dot” between the paired convergent CTCF binding sites indicating the canonical chromatin loop, the cohesin ChIA-Drop data also displayed loop “dots” between cohesin’s loading and anchoring sites, revealing the anticipated chromatin loop formation if cohesin-mediated extrusion is initiated from the loading site and completed at the anchoring site. In comparison, the aggregation plot of CTCF ChIA-Drop data showed negligible ‘cross-shaped’ signals in the center but exhibited the expected strong looping ‘dot’ between the two convergent CTCF sites and the characteristic anchor-based stripes (**Figure 2D**, top middle panel). Similarly, the cohesin ChIA-PET data also displayed the same strong mapping signals of the ‘cross-shaped’ pattern from the loading site as observed in cohesin ChIA-Drop data, while the CTCF ChIA-PET data exhibited the typical anchor-based stripes and dots same as shown in the CTCF ChIA-Drop data (**Figure 2D**, bottom left and middle panels). In addition, the RNAPII ChIA-PET data also showed enriched ‘cross-shaped’ signals in the center at the loading site (**Figure 2D**, bottom right panel) but lacked signals from the loop anchor sites, which is consistent with the current view that cohesin often loads onto DNA at active transcription regions and co-localize with RNAPII.^35,50,75,76^ To rule out potential bias introduced in the ChIP-enriched experiments, we utilized the *in situ* Hi-C data^5^ for the same aggregation analysis and demonstrated the same profile as displayed in the combined views of the cohesin and CTCF plots (**Figure 2D**, top right panel) except for a much broader diagonal line reflecting the high level of random noise resulting from local polymer contacts. Thus, our genome-wide aggregation peak analyses clearly demonstrated the lack of ‘jet’ signals and suggested that cohesin at its loading sites extrudes DNA in a one-sided manner towards either direction.

The one-sided pattern is also apparent as exemplified at individual loci. For instance, on chromosome 4 (Chr4) in a 420 kb segment of a chromatin folding domain defined by CTCF and cohesin ChIA-PET data (**Figure 2E**), 4 distinctive cohesin loading sites within this CTCF loop domain were displayed by ChIP-seq peak signals of NIPBL and associated protein markers. The 2D contact heatmap and browser loop/peak views showed extensive chromatin interactions between cohesin loading sites (L) and anchoring sites (A) as possible loading-to-loading (L-L), loading-to-anchor (L-A), and anchor-to-anchor (A-A) loops. In another example on chromosome 2 (**Figure S2D**), a cohesin-loading block with multiple loading loci resides in the middle of a well-defined CTCF/cohesin loop. Even though the 2D contact heatmap of Hi-C data may seemingly appear as a ‘jet’ signal originating from the cohesin loading block, most likely due to lower mapping resolution (**Figure S3**), the cohesin ChIA-PET data with much higher mapping resolution and specificity clearly revealed distinctive “one-sided” extrusion pattern from individual cohesin-loading sites and observable chromatin contact “dot” as loops between loading sites (L-L loops), in addition to the loading-to-anchor (L-A loops) and anchor-to-anchor (A-A loops).

To elucidate cohesin-mediated chromatin extrusion in single-molecule resolution, we visualized the cohesin ChIA-Drop looping data in browser views, in which enrichment of chromatin fragments aligned to cohesin-loading and -anchoring sites were clearly observable (**Figure 2F**). The ChIA-Drop data in single-molecule resolution could also be viewed to display extrusion trajectory. We sorted the chromatin fragments in the ChIA-Drop data by focusing on one of the cohesin-loading sites and observed that the “one-sided” asymmetric pattern of single-molecule contact events (n=152) was 7-fold higher than the presumed “two-sided” extrusion (n=22) in a symmetric pattern, indicating that the translocation of cohesin from its loading sites are predominantly one-sided, while the symmetric mapping data are most likely mapping noise (**Figure 2G**). An additional example exhibited the similar profiles as presented in **Figure S2E-F**.

The genome-wide statistics of the 3808 cohesin loading sites and the associated 4110 cohesin anchoring sites of the 2055 CTCF/cohesin loops (**Figure 2H**) provided further quantitative measurement in supporting our observations at individual loci. Overall, the counts of the presumed ‘two-sided’ extrusion contacts (median=29) was significantly lower than that of ‘one-sided’ extrusion contacts (median=62) at cohesin-loading sites (**Figure 2H** left panel, p value<1.8e-308). As a likely negative control at the cohesin-loading sites where there were no detectable CTCF binding signals, the CTCF ChIA-Drop counts reflected the background level of signals (two-sided median=18, one-sided median=16, p-value=0.01) (**Figure 2H** right panel; **Figure S2F**), suggesting that the ‘two-sided’ mapping observed in the cohesin data were at the similar level as background noise.

Together, our mapping analyses through aggregation of thousands of well-defined cohesin loading sites, individual examples in bulk cell ensemble and single-molecule mapping trajectory, and genome-wide statistics provided experimental evidence to strongly suggest that the ‘one-sided’ asymmetric model is likely the primary mechanism for cohesin-mediated chromatin loop extrusion from its loading sites in human cells.

### Cohesin anchors at CTCF sites and reels in DNA into loops in a unidirectional manner

It had been generally believed that a cohesin ring complex translocate along the DNA track until it is blocked by CTCF at its binding site.^77–79^ New evidence based on protein crystal structure and functional assays suggested that when a cohesin/NIPBL complex arrives at a CTCF binding site, the N-terminus of CTCF competes with WAPL to interact with cohesin, which would release NIPBL and anchor the cohesin at CTCF site.^80^ In this scenario, the CTCF/cohesin complex would start to reel in the DNA template in a one-sided and unidirectional manner.

To test this model *in vivo* using CTCF- and cohesin-specific mapping data, we analyzed the chromatin contacts derived from both cohesin- and CTCF-enriched ChIA-Drop and ChIA-PET datasets. The chromatin interaction mapping profiles with distinct patterns of stripes and loops as reflected in the 2D contact maps and the browser-based loop/peak views were remarkably similar for both CTCF and cohesin data as exemplified in a 5 Mb segment of Chr3 in GM12878 cells (**Figure 3A-B**). This result is surprising considering that CTCF and cohesin each has very different nature involved in DNA loop making with CTCF binding statically while cohesin extruding progressively. To delve in further, we zoomed in and examined two individual architectural domains within this segment. The first domain (I) encompasses a large loop and two sub-loops displaying multiple CTCF binding peaks and motifs in convergent orientation, while the second domain (II) has a rather simple structure with a dominant large loop as demonstrated in the loop/peak views of the ChIA-PET data, along with ChIP-seq data precisely demarcating the corresponding chromatin features (**Figure 3B** bottom panel). The single molecule mapping in fragment views of the cohesin and CTCF ChIA-Drop datasets displayed the enrichment of contacts at corresponding CTCF loci (**Figure 3C** top panel), and the sorted fragment views based on contact distance (smallest to largest) from CTCF/cohesin anchoring sites revealed a dynamic and continuous trajectory of loop extension in concordance with the CTCF motif orientation at single molecule resolution (**Figure 3C** bottom panel). Additional examples of single molecule resolved extrusion trajectories originated from cohesin loading and anchoring sites are presented in **Figure S4**. Strikingly, the detailed single-molecule mapped looping events in progressive trajectories are almost identical as displayed in both cohesin and CTCF data. Indeed, genome-wide assessment of the cohesin and CTCF ChIA-Drop data are also highly correlated in terms of binding peak intensity and looping frequency (**Figure 3D**).

**Figure 3.**
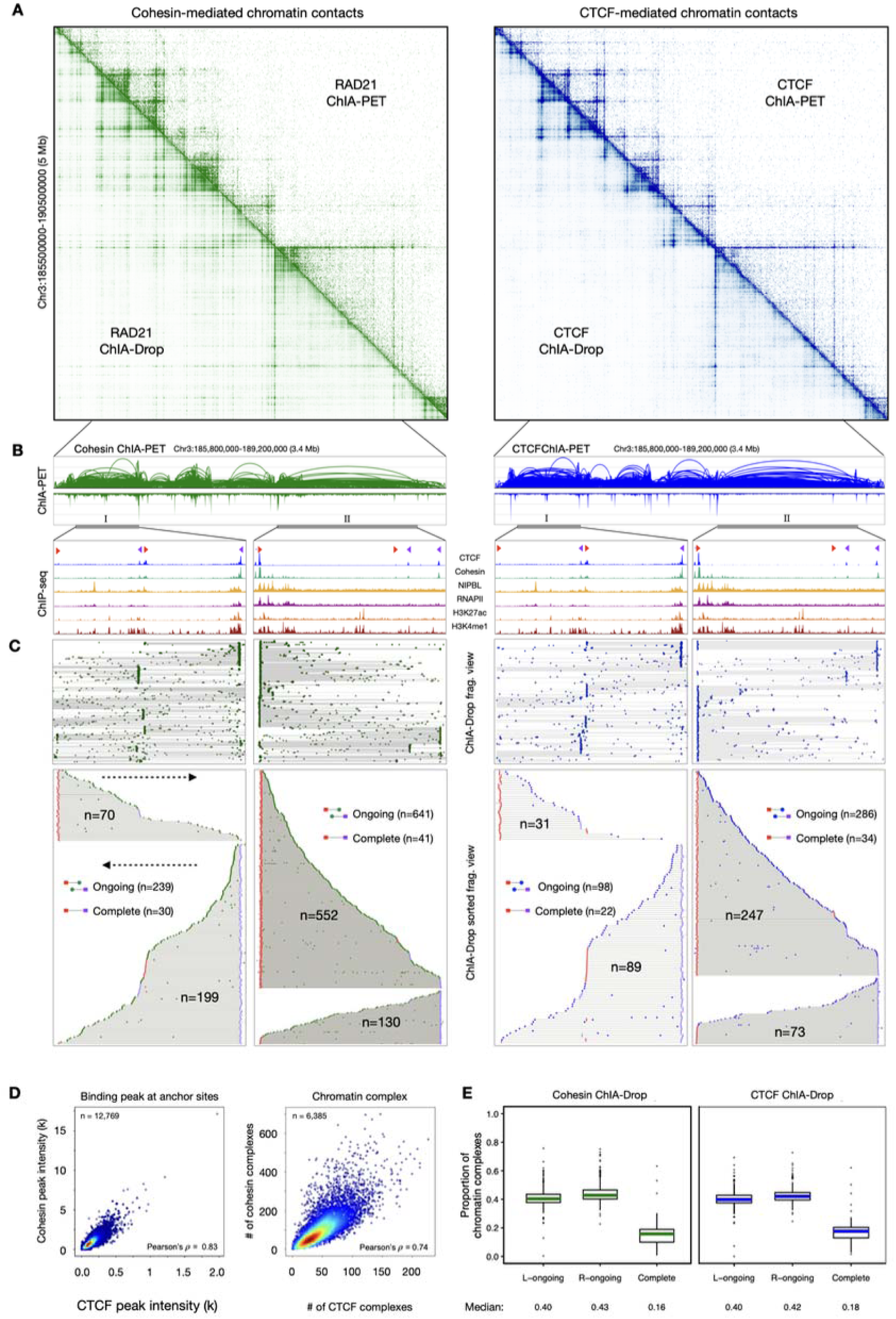
Dynamic looping at CTCF/cohesin anchoring sites. (**A**) An example locus of 2D chromatin contact maps of the RAD21 (green, left) and CTCF (blue, right) ChIA-PET (upper half) and ChIA-Drop (lower half) data showing similar contact profiles. (**B**) Zoom-in browser views of cohesin and CTCF ChIA-PET loops and peaks. Further zoom-in of the marked domains I and II for epigenomic features of ChIP-seq defined cohesin loading and anchoring loci. Position of CTCF binding site motifs are also included (red arrow for forward, purple arrow for reverse orientation). (**C**) ChIA-Drop fragment views (upper) and the sorted fragment views (lower) of chromatin interaction complexes anchored at convergent CTCF motifs and extended in the middle of the looping domain (cohesin green solid dots, CTCF blue solid dots). The directions of the projected trajectories are indicated by dotted arrows. The numbers of ‘ongoing’ and ‘complete’ looping events are provided. (**D**) Scatter plots showing the correlation between CTCF and cohesin ChIA-Drop binding peak intensity at anchors of chromatin loops with convergent CTCF motifs (left) and the numbers of chromatin complexes in CTCF and cohesin ChIA-Drop data at loop anchoring sites (right). (**E**) Boxplots for the proportion of the left anchor originated ongoing (L-ongoing), right anchor ongoing (R-ongoing), and complete looping events. The medians are provided.

It is assumed that a specific looping event in a cultured cell mass would occur with a slightly time-point-off in individual cells; hence, the snapshot of ChIA-Drop single molecule mapping data from a cell population would collectively depict the chronological progression of loop extrusion in a sorting scheme. In this context, as depicted in the sorted views (**Figure 3C**, bottom panel) approximately 10-20% of the chromatin extrusion events are considered as “complete looping” structures, in which the two anchoring points at the convergent pairs of CTCF binding sites are connected. The vast majority (about 80%) are associated with only one of the two CTCF anchors (**Figure 3E**), reflecting a continuous trail of progressing extrusion events (ongoing looping) and implying that cohesin-mediated chromatin looping is a highly dynamic process. Intriguingly, as exemplified in “domain I”, there were also looping events that appeared to bypass CTCF bound and continued to extrude until encountered another CTCF blocker. This phenomenon may reflect the stochastic oscillation of CTCF binding kinetics “on” and “off” DNA at specific loci and moment. When CTCF is “on”, the incoming CTCF/cohesin extrusion complex would be arrested there resulting in a completion of looping, but when CTCF is “off”, the extrusion complex would pass through and continue to extrude.

Together, our results provided compelling mapping evidence that CTCF does not simply function as an insulator passively blocking cohesin extrusion along chromatin *in vivo*. Instead, CTCF actively provide an anchoring point for cohesin and dictate the orientation of extrusion. This notion from our mapping perspective is consistent with previous studies showing that CTCF could stabilize cohesin on the chromatin during extrusion.^80,81^

### Depletion of cohesin and CTCF validates their collaborative efforts in chromatin looping

To validate our observations from GM12878 cells, we exploited the well-established auxin-inducible degron (AID) system^82^ for acute depletion of RAD21 and CTCF to independently measure their effect on loop extrusion starting from cohesin’s loading sites in K562 cells. We first generated high quality NIPBL CUT&Tag binding data along with available ChIP-seq and ChIA-PET data from K562 cells for identification of cohesin loading and associated anchoring sites (**Figure S5A**), and then we produced RAD21 ChIA-PET datasets from the no AID treatment control cells without auxin treatment as wild-type (WT) and RAD21-depleted (RAD21-AID) cells via auxin treatment to analyze if the mapping signals of chromatin “stripes” and contact “dots” stemming from cohesin-loading sites are cohesin-dependent (details in **Methods**; **Figure S5B-C**). Applying the same criteria used in GM12878 cells, we identified higher quality NIPBL binding sites (n=5672) with support of RNAPII, H3K27ac, and H3K4me1 peaks, from which we selected higher confident cohesin-loading sites (n=2681) that are encapsulated in well-defined convergent CTCF loop domains (n=1766) and are far away from CTCF binding sites in K562 cells (see **Methods**). The 2D contact aggregation plots of the RAD21 ChIA-PET data (**Figure 4A**) centered on these cohesin loading sites flanked with associated convergent CTCF sites from control wild type (WT) K562 cells showed the similar 2D contact profile as the GM12878 data (**Figure 2D**), displaying the ‘cross-shaped’ signal from the cohesin loading site in the center and the corresponding loading- and anchoring-based extrusion “stripes” and loop “dots”. Upon RAD21 depletion (RAD21-AID), all contact signals in genome-wide aggregation plot are visibly attenuated (**Figure 4A**), which is particularly noticeable in the loss of signals in the anticipated loading- and anchoring-sites as well as stripes and dots of loading-to-anchoring and anchoring-to-anchoring loops (**Figure 4B**). At two specific loci as shown in Chr8 and Chr10 (**Figure 4C**), distinctive stripes stemming from specific cohesin loading site as demarcated by the peaks of NIPBL and associated protein factors are interconnected forming loop dots between loading sites in control cells; however, after RAD21 depletion (RAD21-AID), those stripes and dots are faded away into the background (**Figure 4C**), which is even more pronounced in the browser-based loops and peaks views (**Figure 4D, S5C**). Our observation of RAD21 depletion in K562 is further supported by similar experimental results in HCT116 cells (**Figure S6**).

**Figure 4.**
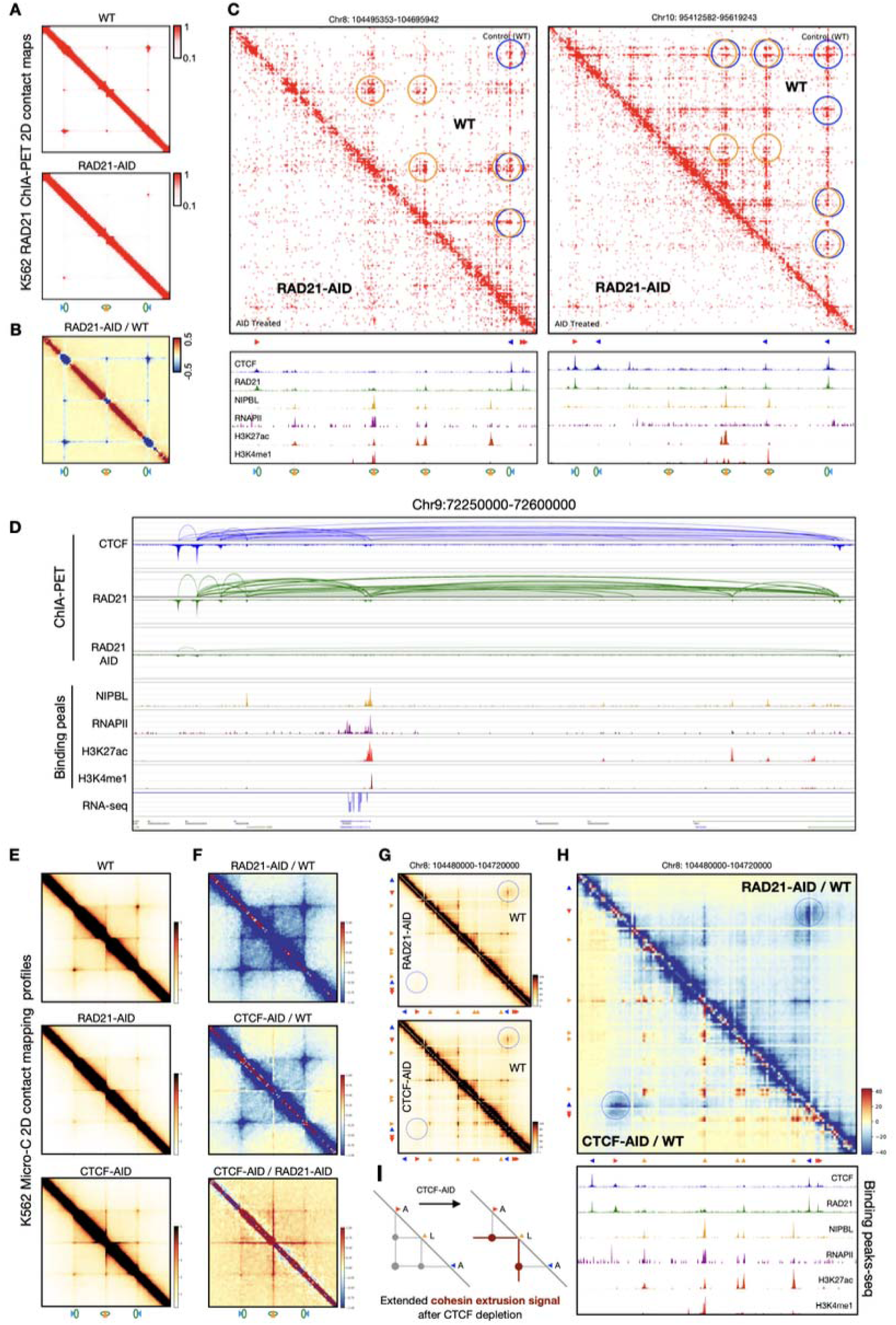
The impacts of acute depletion of RAD21 and CTCF on cohesin-mediated extrusion in K562 cells. (**A**) 2D contact maps of aggregating RAD21 ChIA-PET data from no-AID treatment wild-type control (WT, top) and from RAD21-AID treated cells (RAD21-AID, bottom) centered at cohesin loading sites (n=2681) and flanked with cohesin anchoring sites. (**B**) Differential subtraction view of the RAD21-AID mapping signals minus WT signals. The blue color intensities indicate the reduction of mapping signals in the RAD21-AID treated cells. (**C**) Comparative 2D contact triangle maps of RAD21 ChIA-PET data at two example loci before (WT) and after (RAD21-AID) depleting RAD21 in K562 cells. Corresponding CTCF binding motifs and cohesin loading and anchoring sites identified by ChIP-seq data are presented. The CTCF/cohesin anchor-to-anchor (A-A, blue circles), loading-to-anchor (L-A, orange/blue circles), and loading-to-loading (L-L, orange circles) loops are marked. (**D**) An example browser view of a chromatin loop domain. Browser tracks from the top to bottom are: CTCF ChIA-PET loops/peaks, cohesin ChIA-PET before (RAD21) and after depleting RAD21 (RAD21-AID), binding peaks for NIPBL and associated factors, and RNA-seq data. (**E**) Aggregated 2D contact maps of Micro-C data from no AID treatment WT (top), RAD21-AID treated (middle), and CTCF-AID treated (bottom) centered at cohesin loading sites with flanked anchoring sites. (**F**) Subtractive 2D contact maps of RAD21-AID / WT, CTCF-AID / WT, and CTCF-AID / RAD21-AID. The blue signal intensity indicates the loss of extrusion stripes and loops in perturbed cells; the red signal intensity indicates the gain of extrusion stripes and loops upon CTCF depletion. (**G**) Comparative 2D contact maps of a chromatin domain. Top: RAD21-AID (lower half) vs. WT (upper half). Bottom: CTCF-AID (lower half) vs. WT (upper half). The positions of anchor-to-anchor (A-A) loop dots are indicated by dotted circles. The relative positions of the corresponding CTCF binding motifs (red and blue horizontal pointers) and the cohesin loading sites (yellow upright pointers) are provided. (**H**) Subtractive and comparative 2D contact maps of the same chromatin domain as in **G** but enlarged for more detailed view. The upper half is a subtraction of RAD21-AID minus WT, and the lower half is a subtraction of CTCF-AID minus WT. The positions of anchor-to-anchor (A-A) loops are indicated by dotted circles. The blue signal intensity indicates the loss of extrusion stripes and loops in perturbed cells; the red signal intensity indicates the gain of extrusion stripes and loops upon CTCF depletion. The ChIP-seq peaks used for defining cohesin loading and anchoring sites are presented. (**I**) A schematic elucidating the augmented cohesin extrusion as observed in extended stripes beyond CTCF boundaries and increased contact dots of loading-to-loading loops upon CTCF depletion.

To further investigate the interplays between cohesin and CTCF, we depleted a cohesin subunit RAD21 and CTCF in K562 cells, respectively, and followed by Micro-C^83^ analysis for deep coverage and higher mapping resolution in the control and auxin-treated cells (details in **Methods**; **Figure S5D-E**). While the RAD21 depletion attenuated all loading- and anchoring-associated signals including extrusion stripes and contact dots (**Figure 4E**) same as observed in RAD21 ChIA-PET analysis (**Figure 4A**), the CTCF depletion appeared to diminish only the CTCF/cohesin anchoring-site-associated signals of extrusion stripes and contact dots (A-A loops) but not affect cohesin loading-site associated signals (**Figure 4E**). Specifically, in the CTCF-AID Micro-C data the chromatin extrusion signals originating from cohesin loading sites appeared not only elevated but also extended beyond the adjacent CTCF boundaries that normally defines the boundaries of convergent CTCF loops (**Figure 4F**). As exemplified at specific loci (**Figure 4G, S7**), while the RAD21-depletion demolished all stripe and dot signals associated with cohesin loading and anchoring loci, the CTCF depletion diminished only the mapping signals associated with cohesin anchoring loci where there are strong CTCF binding. Strikingly, the differential/subtraction analysis comparing CTCF depletion with RAD21-depletion data over the wildtype (WT) not only validated the CTCF-dependency of anchor-based looping but also, unexpectedly, highlighted the substantial elevation of cohesin loading specific stripes and particularly the chromatin contact dots of loading-to-loading (L-L) loops that are closely associated with the NIPBL presence and are CTCF-independent (**Figure 4H** top panel, **4F** bottom panel). While the loss of CTCF is expected to allow cohesin extrusion extending in further length, the elevation of L-L loop signals is unexpected, which hint that when CTCF is diminished, the system may respond to have more cohesin activities to compensate the loss of CTCF (**Figure 4I**, more in Discussion).

### Loop extrusion from cohesin loading, anchoring, and to completion

From the results of our analyses and by incorporating recent reports,^28,43,47,80^ it becomes evident that the entire process of cohesin-mediated extrusion for chromatin loop formation likely involves two continuous phases: the initial cohesin extrusion from its loading site and the second extrusion at its anchoring site to complete chromatin looping (**Figure 5A**). Our proposed model is as follows.

**Figure 5.**
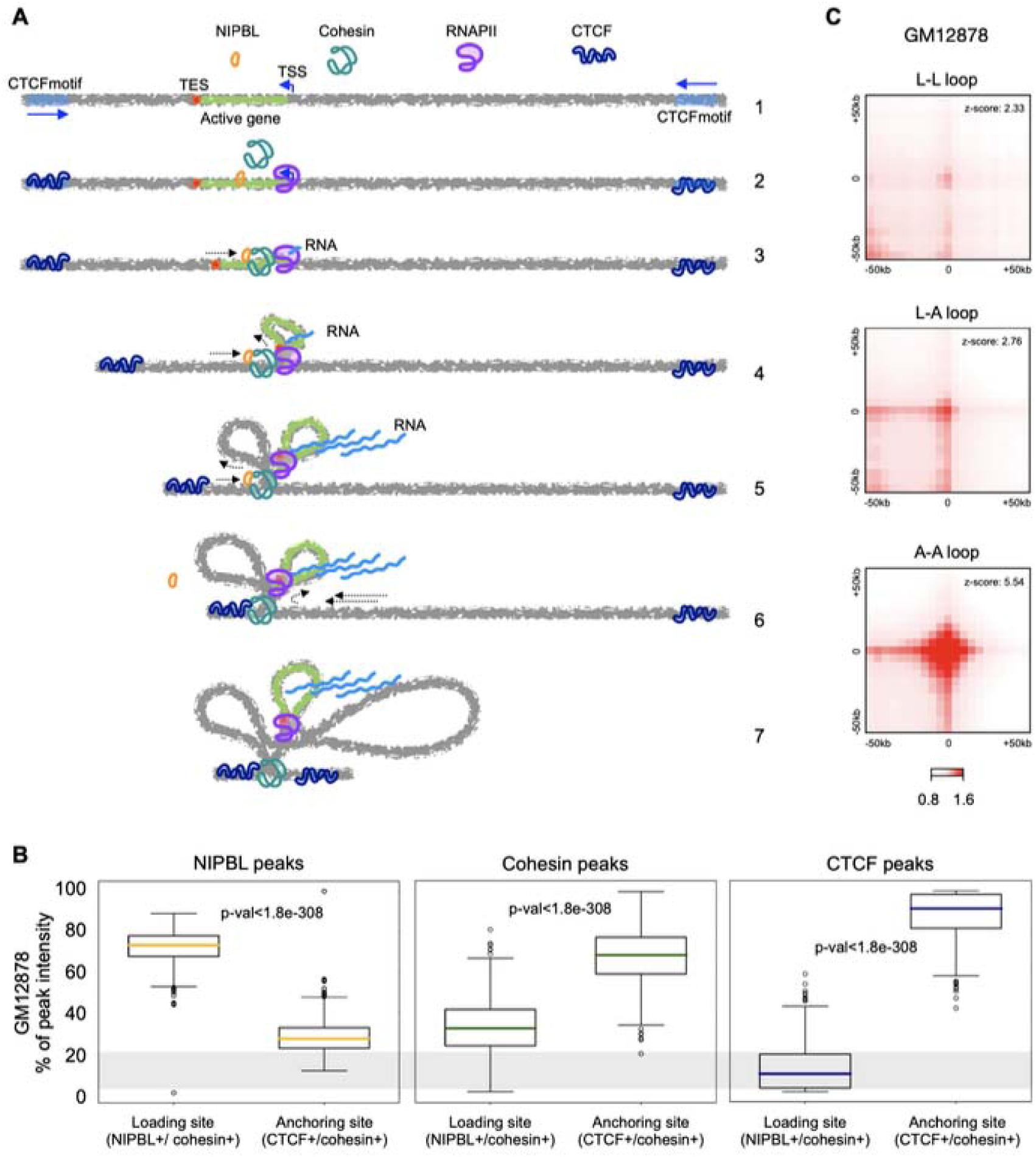
A one-sided and two-mode model of cohesin loop extrusion. (**A**) The model. Step 1: Convergent CTCF looping domain usually contains active transcription regions and NIPBL binding sites. 2: Cohesin loads at NIPBL binding site, which often overlaps with active transcription region and occupies by RNAPII. 3: After loading, cohesin interacts with NIPBL to form NIPBL/cohesin complex, which extrude DNA together with RNAPII to establish transcriptional loop. 4: The NIPBL/cohesin complex together with RNAPII extrude DNA to establish transcriptional loops. 5: After transcription, NIPBL/cohesin will continue extrude until encountering the N-terminal of a CTCF at its binding site with a motif in a opposite orientation. 6: The CTCF and cohesin interaction will release NIPBL, and the CTCF/cohesin protein complex will somehow switch extrusion direction and start to reel in DNA backward. 7: CTCF/cohesin will continue to reel in DNA until arriving to another CTCF in the context of convergent CTCF loop domain, thus to complete the loop formation. (**B**) Genome-wide assessment of binding peak intensity at the cohesin loading and anchoring sites by NIPBL (left), cohesin (middle), and CTCF (right) in GM12878 cells. More than 70% of the NIPBL binding signals are at the cohesin loading sites, and less than 30% associated with anchoring sites. On the contrary, while cohesin loads onto DNA with NIPBL, more cohesin signals are associated with CTCF at anchoring sites, indicating a nature of cohesin, staring with NIPBL for loading and ended up with CTCF for anchoring. The CTCF peak data here provides a reference that more than 90% of the CTCF binding signals are associated with cohesin anchoring sites and the least at cohesin loading sites, illustrating CTCF’s exclusive specificity as cohesin’s anchoring point. **(C).** Aggregation plots of chromatin loops between loading-to-loading (L-L), loading-to-anchoring (L-A), and anchoring-to-anchoring (A-A) loops in GM12878 cell line. The APA z-score of the A-A loop is more than 2-fold, indicating that A-A loop is the predominant looping function in establishing 3D genome structures.

In the first phase, cohesin loading is highly specific at NIPBL binding sites, and most of NIPBL binding sites locate in transcriptionally active regions distal to CTCF binding sites. The NIPBL/cohesin complex starts to extrude DNA in a “one-sided” manner without direction preference; however, if there are active gene promoters co-localized with cohesin loading sites, cohesin extrusion generally follows the same direction as gene transcription.^35^ At the end of the first phase, the extruding cohesin encounters CTCF at its N-terminus region, which would result in a transient loading-to-anchoring (L-A) loop. NIPBL in the NIPBL/cohesin complex is replaced by CTCF, and the cohesin is then anchored at the CTCF binding site in the form of a CTCF/cohesin complex.^80^ This model is consistent with our mapping results showing that the relative cohesin binding signals at its loading sites with NIPBL are much weaker than those at its anchoring sites coupled with CTCF (**Figure 2A-B**, **Figure 5B**), indicating that although most of cohesin are not co-localized with CTCF but eventually accumulated at CTCF sites.

In the second phase after cohesin is anchored with CTCF, there are possible two scenarios. One popular and widely perceived scenario would be that cohesin stops extrusion and stalls there by the insulator function of CTCF same as the “two-sided” model suggested. However, this scenario could not explain why most strong chromatin interactions are anchor-to-anchor (A-A) loops with convergent CTCF sites. An alternative scenario is that cohesin would continue to extrude at the CTCF anchoring site but in a different manner. Our aggregation analysis of mapping data showed that the loading-initiated chromatin looping intensity (L-L and L-A loops) are about 2-fold less than the anchoring-based (A-A) looping as measured by APA analysis in Z-score (**Figure 5C**). Our perturbation experiments further revealed that the loading site-associated L-L and L-A loops are cohesin-dependent but CTCF-independent, while the anchor-associated A-A looping are sensitive to both CTCF- and cohesin-depletion (**Figure 4E-F**). Thus, our results functionally uncovered the existence of two interconnected extrusion mechanisms mediated by NIPBL/cohesin responsible for extrusion from loading to anchoring and CTCF/cohesin for extrusion from anchor to anchor, respectively. These two extrusion processes are tightly connected at the junction point of CTCF binding sites (**Figure 5A**). If cohesin loading sites are proximal to CTCF sites, it may immediately start the CTCF/cohesin-mediated extrusion phase (**Figure S8A**). Our aggregation data also suggest that the CTCF/cohesin-mediated extrusion is the predominant function in chromatin loop formation likely responsible for establishing all convergent CTCF loops (**Figure 5C**), which are the primary chromatin folding constituents in TAD and other higher-order structures.

Our results above prompted a critical question—namely, if it is possible for a cohesin to change its extrusion direction *in vivo* after integrating with CTCF? Several recent *in vitro* experiments have provided insightful structural evidence,^84,85^ suggesting that human cohesin can switch its extrusion in a opposite direction. Therefore, it is likely in vivo that CTCF/cohesin could switch its direction backward to complete chromatin looping between two convergent CTCF sites.

### In silico simulations recapitulate the one-sided and two-mode model of cohesin extrusion

To validate our experimental observations and the proposed one-sided two-mode of cohesin extrusion mechanism, we developed a data-driven computational simulation program (see **Methods**) by modifying the widely-used DNA polymer algorithm ^24,87,88^ with our experimentally obtained parameters as inputs (see **Methods**), and performed *in silico* simulations that model the behavior of DNA polymers *in vivo*. In particular, we compared two possible models: 1) the one-sided two-mode model proposed in this work (**Figure 5A**), in which after loaded at NIPBL sites, cohesin slowly moves toward one of the CTCF anchor sites in a one-sided manner and subsequently switches its direction once encountering CTCF and starts to reel in DNA backward actively in an ATP-dependent, high-force manner until reaching the other convergent CTCF motif; and 2) the widely postulated two-sided one-mode model, wherein cohesin extrudes DNA at NIPBL sites symmetrically in both directions, forming a chromatin loop when it reaches the convergent CTCF sites ^57^. We investigated the feasibility, efficiency, and accuracy of these two models.

As a first step, we examined which of the two models best recapitulates the experimental observations. An 800 kb region on chromosome 4 in K562 cells encompasses multiple CTCF motifs that coincide with CTCF and cohesin binding peaks, representing potential CTCF anchoring regions, and a strong cohesin loading site in the middle as characterized by high quality ChIP-seq peaks of NIPBL, RNAPII, H3K27ac, and H3K4me1 (**Figure 6A**, ChIP-seq/CUT&Tag). The 2D Micro-C contact map of this region shows three main chromatin domain structures: a small loop (dot) with a left stripe on the top-left, a medium-sized left stripe with a cross-shaped signal originating from the loading site in the middle (indicated by an arrow), and a combination of stripes, loops, and domains in the bottom-right corner (**Figure 6A**, Micro-C).

**Figure 6.**
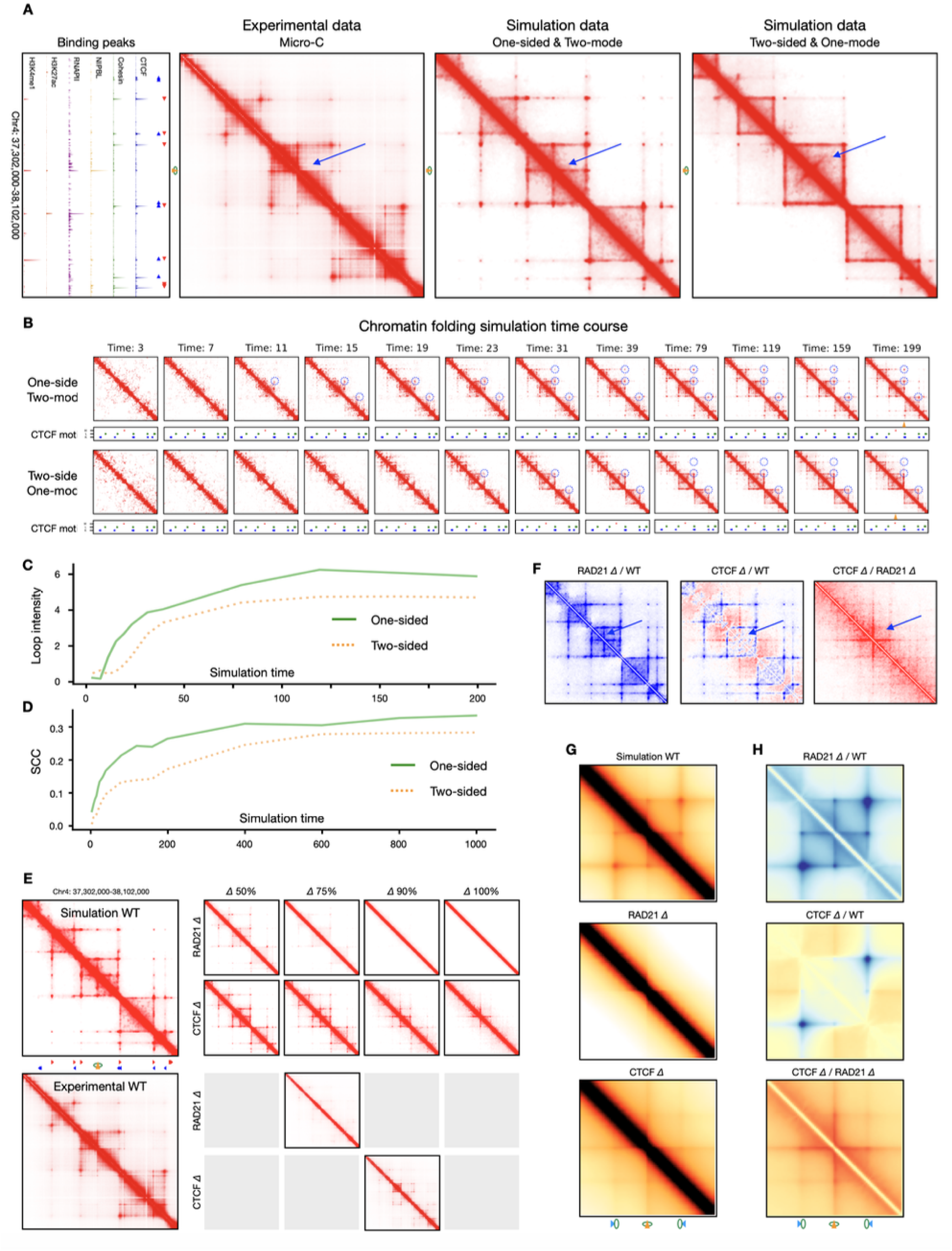
Computational simulations of cohesin-mediated loop extrusion. (**A**) Binding peaks of ChIP-Seq and CUT&Tag data along with CTCF motifs (red and blue triangles) used for defining cohesin loading and anchoring sites in a representative genomic region are presented (left). Three 2D contact maps for this genomic region, the experimental data of Micro-C contact map (left), simulated contact map generated by the “One-sided & Two-mode” model (middle), and the simulated contact map “Two-sided & One-mode” model (right). The major cohesin loading site in middle of the chromatin domain is indicated by an orange triangle (NIPBL site) overlapped with green circle (cohesin ring). The characteristic extrusion patterns, the “cross-shaped” stripes in the one-sided model and the perpendicular jet-like stripe in the two-sided model—are highlighted by blue arrows. (**B**) Time-course series of simulated contact maps comparing the efficiency of loop formation between the two models. Blue circles highlight representative chromatin loops. The “One-sided & Two-mode” model forms loops more rapidly. (**C**) Quantification of anchor-to-anchor (A-A) loop intensity over simulation time for the two models. The loop intensity in the “One-sided” model is stronger than that in the “two-sided” model at all time points. (**D**) Comparison of pattern similarity between simulated and experimental contact maps over time, measured by the HiCRep stratum-adjusted correlation coefficient (SCC). The “One-sided” model shows higher similarity to Micro-C data than the “Two-sided” model. (**E**) Top, simulated 2D contact maps generated by the “One-sided & Two-mode” model under wild-type conditions and *in silico* perturbation with increasing levels of CTCF or RAD21 depletion (Δ50%, 75%, Δ 90%, and Δ100%). Bottom, Corresponding Micro-C 2D contact maps under matched in silico perturbation conditions. Grey squares indicate no experimental data at the matched *in silico* efficiency. (**F**) Differential subtraction 2D contact maps of *in silico* perturbation of RAD21 depletion (Δ100%) over wildtype (RAD21Δ / WT, left), CTCF depletion (Δ100%) over wildtype (CTCFΔ / WT, middle), and CTCF depletion (Δ100%) over RAD21 depletion (Δ100%) (CTCFΔ / RAD21Δ). The “cross-shaped” stripes are indicated by blue arrows. Notably, the “cross-shaped” stripe signals showed extended and increased signals after_CTCF depletion (Δ). (**G**) Aggregation of simulated 2D contact maps from 200 genomic regions centered on cohesin loading sites and flanked by convergent CTCF anchor sites, generated using the “One-sided & Two-mode” model. Aggregation plots for wild-type (WT, top), 100% RAD21 depletion (RAD21Δ, middle), and 100% CTCF depletion (CTCFΔ, bottom) are shown. The corresponding cohesin loading sites and anchoring sites are indicated. (**H**) Differential subtraction 2D contact maps of aggregated signals between conditions: RAD21Δ / WT (top), CTCFΔ / WT (middle), and CTCFΔ / RAD21Δ (bottom).

The simulation of our proposed one-sided two-mode model closely recapitulates the patterns observed in the Micro-C data, notably the cross-shaped signal at the loading site, the stripe signals extended from the anchoring sites, and the contact dots for A-A loops (**Figure 6A**, One-sided & Two-mode). By contrast, simulation of the two-sided and one-mode of extrusion model produced a ‘jet-like’ or ‘fountain-like’ structure at the loading site (**Figure 6A**, Two-sided & One-mode) as expected but contradict the cross-shaped signal observed in experimental data (**Figure 6A**, Micro-C). Moreover, the two-sided stripes appear as solid square boxes, and the model generated several additional structures that are not visible in the Micro-C data. These visual inspections indicate that our proposed one-sided two-mode model can reproduce the observed data more accurately than the two-sided one-mode model.

The simulations shown in **Figure 6A** were at timepoint 999 (each time unit equals 10 simulation steps; see **Methods**), which allows sufficient time for cohesin to extrude and saturate chromatin contacts. We next tested which of the two models is more efficient in forming chromatin structures, especially the convergent CTCF loops. To this end, we reasoned that if one of the major goals of extrusion in vivo is to establish the CTCF anchored chromatin loops, then the simulation timepoint can be used to serve as a proxy for efficiency: the shorter time required to reach the observed contact loops, the more efficient the model. As a result, the one-sided model began forming the first observable contact dot of a convergent CTCF loop as early as at timepoint 11, while the contact dot of the same loop in the two-sided extrusion model emerges as late as timepoint 23 (**Figure 6B**, blue dotted circles). The other loops in this domain also appeared much sooner in the one-sided model than in the two-sided model. These visual patterns are validated by quantifying the loop intensity over the simulation time course (**Figure 6C**). We also quantified the similarity between the simulated contact maps and the experimental Micro-C data including features of loops and stripes. Specifically, we computed the stratum-adjusted correlation coefficient (SCC) using HiCRep,^89^ where the higher SCC value (1 to 0) the higher similarity. Consistent with the visual patterns in **Figure 6B**, the one-sided model consistently showed higher SCC values at every simulation timepoint than the two-sided model, and it also plateaued much earlier (around simulation time 400) than the two-sided model (around simulation time 600) (**Figure 6D**). Together, these results indicate that the one-sided two-mode model is more feasible, accurate, and efficient than the widely believed two-sided one mode extrusion model.

### Polymer simulations reproduce the effects of cohesin and CTCF depletion

Having demonstrated that the one-sided two-mode extrusion model could better recapitulate the experimental data, we next applied our simulation algorithm to mimic the effects of cohesin and CTCF depletion on chromatin interactions via in silico perturbation analysis (see **Methods**).

One advantage of simulation over experimental approaches is its ability to precisely control the amount of effective CTCF and cohesin quantitatively via *in silico* perturbation. We therefore simulated the depletion of CTCF and cohesin at various levels from 0% (no depletion control as WT) to 100% (complete depletion). Through a in silico titration of perturbations, we observed progressive changes in simulated contact maps (**Figure 6E** upper panel). Interestingly, RAD21-depletion at 75% already disrupted most of the looping structures and completely abolished all looping signals at 90% depletion, reflecting the importance of cohesin in chromatin looping. In contrast, 75% of CTCF depletion appeared to have only minimal impact to looping features, and the extrusion stripes from the cohesin loading site are retained even when CTCF is completely removed (100% depletion), exactly matching with our experimental Micro-C data (**Figure 4E-F**). Referencing the results of in silico perturbation titration, our RAD21-AID experiment data probably reflected a depletion efficiency at 75% to 85%, while our CTCF-AID experiment closely matches the simulation result of 90% CTCF in silico depletion (**Figure 6E** lower panel).

To compare the simulation results, we plotted the pairwise differences in 2D contact signals between conditions. As a result, the RAD21 in silico depletion relative to WT showed substantial loss of both loading- and anchoring-associated extrusion stripes and loops, whereas the CTCF in silico depletion relative to WT diminished only the anchor-associated extrusion signals but retained loading stripes (**Figure 6F**, left and middle; **Figure S9A-B**). Interestingly, direct comparing the *in silico* depletion of CTCF with RAD21 clearly revealed remarkable elevation and extension of cohesin specific extrusion signals originated from cohesin loading site (**Figure 6F**, right)—all consistent with our Micro-C mapping data (**Figure 4F**) and, thus, provided an orthogonal confirmation of our experimental observation.

To further evaluate these effects at a genome-wide scale, we performed simulations for all regions centered on loading sites in K562 cells and conducted the same aggregation analysis as in the previous sections. The aggregated 2D contact maps (**Figure 6G-H, S9C**) exhibited similar patterns as simulated for individual region (**Figure 6E**), which is also consistent with our experimental observations (**Figure 4E-F**). Collectively, these analyses of subtractive in silico perturbation demonstrate that the one-sided two-mode extrusion model faithfully reproduces the chromatin interaction changes observed upon cohesin and CTCF depletion. More importantly, the simulation results provide *in silico* support to our seemingly surprising experimental results: removing CTCF elevated loading-based stripe signals and loading-associated loops.

## DISCUSSION

Cohesin has been in an active area of research due to its importance in forming chromatin loops, regulating gene expression, and establishing 3D genome organization. In addition to the substantial progress being made to understand the mechanism of cohesin-mediated loop extrusion through *in vitro* studies, considerable efforts have been invested in understanding the mechanisms *in vivo* through depletion of various protein factors-CTCF, RAD21, WAPL, NIPBL-to measure their effects on 3D genome organization. ^31,33,41,44,45,51,62^ However, most of these studies focused on CTCF-based chromatin loops without addressing extrusion derived from defined cohesin loading sites or lacked necessary specificity and resolution. In this study, we leveraged high-resolution and high-specificity mapping approaches coupled with perturbation of protein factors and centered on well-defined cohesin loading and flanked anchoring loci in the context of canonical convergent CTCF loops to delineate the intricate interplays of cohesin with NIPBL and with CTCF during chromatin loop formation in human cells. We provide sufficient *in vivo* evidence to deduce that cohesin extrudes DNA predominantly in a ‘one-sided’ unidirectional manner from its initial loading sites and at its anchoring site. By incorporating recent *in vitro* evidence with our *in vivo* mapping data, we postulate a working hypothesis, in which cohesin with NIPBL extrudes DNA asymmetrically from its loading sites; once anchored with CTCF, it switches its extrusion direction backward and reels in DNA unidirectionally until complete the convergent CTCF loop in a highly dynamic fashion. This one-sided two-mode extrusion mechanism challenges the current notion that cohesin extrudes DNA in a two-sided bidirectional manner ^57,90^ and provide a comprehensive view of cohesin-mediated extrusion from its initial loading sites and completion in the context of convergent CTCF loops.

One of the difficulties in studying cohesin-mediated chromatin extrusion *in vivo* has been the lack of trustworthy locations in which cohesin precisely loads onto DNA as the starting point of extrusion. Although it has been reported in literatures that cohesin loads onto DNA where NIPBL is bound,^52^ NIPBL interacts with DNA not through sequence-specific loci with recognizable motifs but rather generally at transcription active regions.^35^ A technical hurdle was the lack of reliable antibody against NIPBL for quality ChIP-seq measurement until recently. The high-quality NIPBL binding data produced in this study enabled us to reliably identify high confidence cohesin loading sites genome-wide, and our comprehensive mapping approaches allowed us to examine specific extrusion trajectories that originated directly from individual cohesin loading and anchoring sites with unprecedented resolution and specificity.

When centered on thousands of well-defined cohesin loading sites in multiple human cell lines, all of our mapping data including ChIA-Drop, Hi-C, Micro-C, and ChIA-PET displayed the characteristic “cross-shaped” stripes in aggregation, and our ChIA-Drop data showed single-molecule-resolved “one-sided” trajectories at individual loading loci, without any observable jet-like or fountain-like signals.^61,62^ Additionally, the observed “cross-shaped” patterns in aggregation and at individual cohesin loading loci are specifically cohesin-dependent and resistant to CTCF depletion, which further validated our observations. Collectively, we conclude that ‘one-sided’ asymmetric extrusion from the loading sites towards either direction is the predominant action in human cell. We postulate that the previously reported jets presumably formed by “two-sided” extrusion might only exist in quiescent cells^60^ and that fountains are likely artifacts due to depleted functions in the perturbed mouse cells.^62^ Characteristically, all reported jets and fountains are originated from large regions enriched for enhancers over tens of thousands kb in size, instead of at individual cohesin loading loci with peak-size resolution (∼500 bp), and they appeared to be diffused in extremely large space in ribbon shape usually over megabases, rather than the commonly observed chromatin extrusion stripes in much narrowed shapes as observed in canonical convergent CTCF loops average in 200 kb size.^65^ It might be possible that some of the reported fountain-like structures seen in zebrafish^64^ and C. elegans^63^ are for other nuclear functions, such as DNA replication coupled fountains^66^, rather than for cohesin-mediated loop extrusion.

In addition to the canonical anchoring-to-anchoring (A-A) loops, our mapping analysis also detected distinctive extrusion stripes originated from cohesin loading sites and recognizable contact “dots” between adjacent loading sites as loading-to-loading (L-L) loops in 2D contact heatmaps. Similarly, we also detected loading-to-anchoring (L-A) loops in our mapping data. These L-L and L-A loops were previously underappreciated most likely due to the lack of reliable NIPBL/cohesin loading information. As expected, the extrusion stripes originated from cohesin loading sites extended beyond CTCF boundaries when CTCF is removed experimentally or computationally. Unexpectedly, the cohesin-mediated extrusion signals associated with L-L and L-A loops were observably elevated upon depleting CTCF. We speculate that the acute depletion of CTCF might trigger some nuclear responses. If establishing CTCF loops is a major goal in 3D genome folding with cohesin and CTCF tightly coordinating this process, the removal of CTCF would somehow trigger an alert to compensate for the loss of CTCF by increasing cohesin extrusion activity. This type of compensation mechanism is akin to the increased interactions between super-enhancers ^31^ and increased RNAPII activity upon RAD21 depletion.^35^ Additional ChIP-seq experiments probing RAD21, RNAPII, and other transcriptionally active marks after depleting CTCF could shed light on the functional identities of L-L and L-A loops, and the regulatory mechanism that balance the process of cohesin-mediated DNA loop extrusion.

When we first learned that the cohesin extrudes DNA from its loading sites in a “one-sided” manner and most cohesin loading sites are distal to CTCF sites while major looping activities are between convergent CTCF sites, we realized a conflicting situation: cohesin extrusion from loading to anchoring (L-A) and from anchor to anchor (A-A) in the context of a convergent CTCF looping domain would have a directional conflict. Even if the L-A extrusion occurs before the A-A extrusion, there must be some transition of directional switch from L-A extrusion to A-A extrusion. We have been struggling to reconcile these seemingly contradicting acts into a harmony as current mapping technologies for *in vivo* study would not resolve this problem. Recently, several *in vitro* studies have provided insightful single molecular evidence to suggest that cohesin could switch extrusion direction and that direction switches strongly correlated with the turnover of NIPBL.^48,55^ Specifically, when human cohesin interacting with CTCF at its YDF motif in NTR, it could modulate cohesin subunit STAG1’s DNA affinity and trigger cohesin to switch its extrusion in a opposite direction to reel in DNA into loops unidirectionally.^85^ Furthermore, cohesin is shown to have two action modes: head-hinge movement that resists forces up to 1 pN driven by random diffusive motion, and the head-head movement that resists forces up to 15 pN via ATP ^54^. These two modes of actions in low- and high-energy consumption could fit well with our experimentally observed two phases of loading to anchor extrusion in low-energy and the A-A extrusion in high-energy modes. Thus, the latest *in vitro* studies provided key pieces of evidence to support our working hypothesis of a two-phased cohesin extrusion model *in vivo*. We envision cohesin to extrude DNA from its loading site to the CTCF anchoring site via head-hinge movement with lower energy consumption; and once coupled with CTCF, cohesin can switch its extrusion direction backward and actively reel in DNA via head-head movement in a higher energy consumption mode until encountering at the other end of a chromatin looping domain. This model is concordant with the strong anchor-to-anchor loops and weak loading-to-anchor loops (**Figure 5C, S8**), and the correlation of anchor-based stripe strengths in an ATP-dependent manner.^58^

Furthermore, connecting with our previous report on the interplays between cohesin and RNA polymerase II in regulating chromatin interactions and gene transcription^35^ leads to an emergence of an integrated perspective for the interconnected multifaceted functions of cohesin in transcription regulation and in chromatin loop formation. At its loading sites in active transcription regions, cohesin translocate along chromatin track together with RNAPII to establish transcription loops in both short-range for constitutive genes and long-range for cell-type-specific genes.^35^ After arriving at its anchoring site, cohesin interacts with CTCF and begin to robustly reel in DNA to make architectural chromatin loops. As suggested in this view, cohesin may fulfill its function in gene transcription before its role for chromatin looping chronologically, potentially a new property of cohesin that had not been previously realized. We anticipate that, as advanced perturbation tools and engineered reagents become available, various aspects of cohesin’s roles *in vivo* can be thoroughly tested further in the near future.

## RESOURCE AVAILABILITY

### Lead contact

Further information and requests for resources and reagents should be directed to, and will be fulfilled by the Lead Contact, Yijun Ruan.

### Materials availability

All unique/stable reagents generated in this study are available from the lead contact without restriction.

### Data and code availability

- All genomics data generated and used in this study have been deposited in GEO and are publicly available as of the date of publication. Additional details are provided in the Key Resources Table in STAR METHODS.
- All original code is publicly available at GitHub:

https://github.com/XiaoTaoWang/cohesin-extrusion-analysis.

- The simulation framework used in this study has been deposited at zenodo: https://doi.org/10.5281/zenodo.19093829.
- Any additional information required to reanalyze the data of reported in this paper is available from the lead contact upon request.

## ACKNOWLEDGEMENTS

This study was supported by National Natural Science Foundation of China (32250710678 to Y.R., 32470698 to X.W., and 32400426 to H.C.), and National Human Genome Research Institute (R00-HG011542 to M.K.). The authors acknowledge Xiaoan Ruan, Meizhen Zheng, and Simon Tian for preliminary ChIA-Drop data generation and analyses. We thank Dr. Jiazhi Hu for his gifts of human K562 RAD21-mAID-mClover cell line and K562 CTCF-mAID-mClover cell line created and well characterized by Dr. Jiazhi Hu laboratory.

## AUTHOR CONTRIBUTIONS

Y.R. conceptualized the project. Y.R., M.K., P.W. designed experiments. P.W., L.M. and H.C., generated mapping data. M.K. and X.W. leaded the data analysis efforts with Y.L., L.J., J.H., and T. Y performed computational analyses. X.W. and L.J. developed the data-driven assisted computational simulation algorithm and performed *in silico* perturbation analysis. Y.R. and P.W. wrote the manuscript with major inputs from M.K. and X.W. All authors read, commented, and approved the manuscript.

## Supplementary figures and figure legend

**Figure S1.**
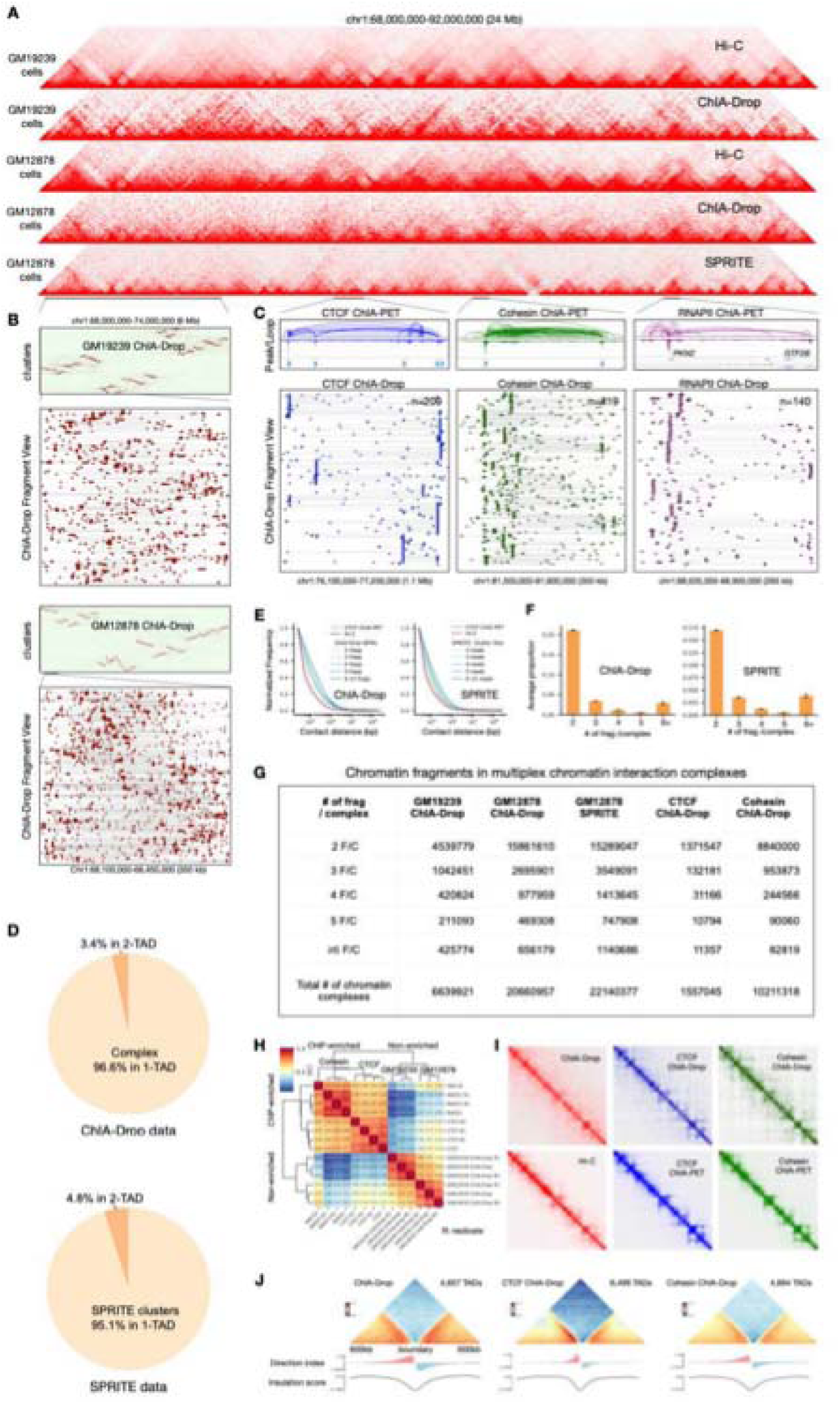
ChIA-Drop data quality assessment (related to Figure 1). **(A)** 2D contact maps at chr1:68,000,000-92,000,000 (24 Mb) in comparison of ChIA-Drop data with Hi-C and SPRITE data derived from GM19239 and GM12878 cells. **(B)** Cluster views (binned) revealing frequently interacting domains and zoomed-in fragment views showing detailed chromatin domains by ChIA-Drop data in GM12878 and GM19239 at chr1:68,000,000-74,000,000 (6 Mb), where each row of dots and a connecting line represents a putative chromatin complex with ≥ 2 interacting fragments. **(C)** ChIP-enriched CTCF (blue), cohesin (green), and RNAPII (purple) ChIA-Drop data are shown with chromatin complexes (bottom panels). Corresponding ChIA-PET loops and peaks at the same loci are included as references (top panels). CTCF binding motifs in CTCF and cohesin data tracks are marked with light blue arrows indicating the binding motif orientation. RNAPII-enriched chromatin complexes are centered around the promoters of active genes PKN2 and GTF2B. **(D)** Proportion of ChIA-Drop and SPRITE complexes spanning in one TAD or two TADs in GM12878 cells. **(E**) Normalized contact frequency and pairwise contact distances of chromatin interactions in ChIA-Drop and SPRITE data broken down by the numbers of fragments per chromatin complex (light green to dark blue) with respect to the CTCF ChIA-PET (dotted red) and Hi-C (dashed red) data as references. **(F)** Bar chart for the numbers of fragments per chromatin complex in ChIA-Drop (left) and SPRITE (right) data. **(G)** Tabulation of fragment numbers per complex (F/C) in non-enriched datasets (ChIA-Drop and SPRITE) and ChIP-enriched datasets (CTCF and Cohesin ChIA-Drop). **(H)** Stratum-adjusted correlation coefficients (SCC) between all datasets of ChIA-Drop and ChIP-enriched CTCF, RAD21 and SMC1A ChIA-Drop experiments and their replicates. The matrix of SCC is clustered via hierarchical clustering. R1, R2, and R3 denote three replicates of a given experiment, respectively. **(I)** Example views of 2D contact matrices of pairwise interactions with 25 kb resolution in non-enriched ChIA-Drop, ChIP-enriched ChIA-Drop data for CTCF or Cohesin (first row), and Hi-C, ChIP-enriched ChIA-PET data for CTCF or cohesin (second row) at chr2:24,000,000-32,000,000 (8 Mb). **(J)** TAD directionality and insulation score called for each of the ChIA-Drop datasets.

**Figure S2.**
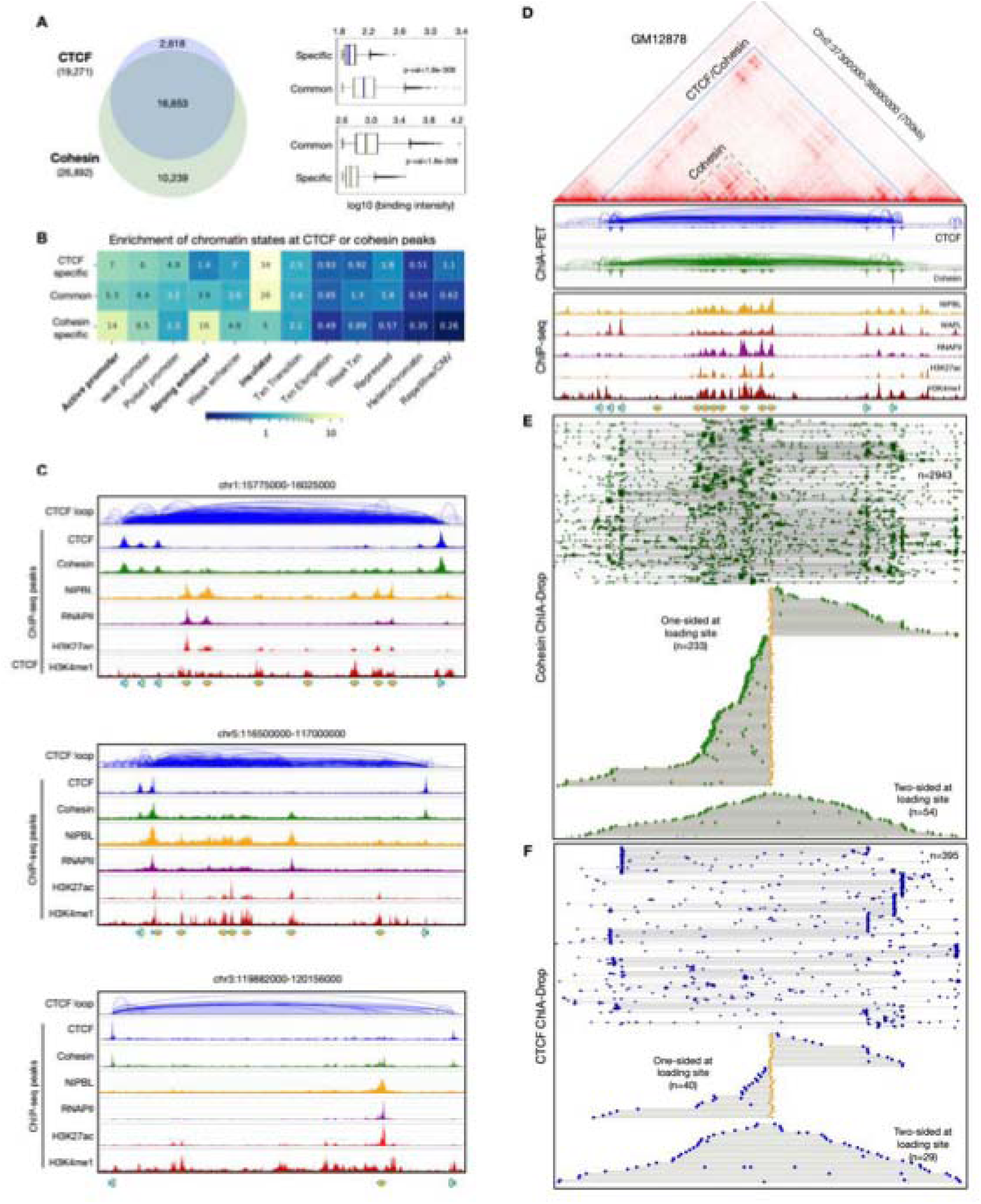
Identification and characterization of cohesin loading and anchoring sites in GM12878 cells (related to Figure 2). **(A)** A Venn diagram of CTCF and cohesin ChIA-Drop peaks (left) with box plots of binding intensity (right) for categories of CTCF-specific, CTCF/cohesin common, and cohesin-specific binding peaks. **(B)** Chromatin states for the three categories of CTCF-cohesin binding loci are characterized *via* ChromHMM annotations. Scores for each ChromHMM feature indicate the fold-change enrichment over 100 random regions. CTCF specific peaks and common peaks between CTCF and cohesin are enriched in insulator marks (bolded), whereas Cohesin specific peaks are enriched in active promoter and strong enhancer marks (bolded). **(C)** Three examples of browser views showing CTCF/Cohesin anchoring sites and cohesin loading sites defined by both CTCF ChIA-PET data and related CTCF, cohesin, NIPBL, RNAPII, H3K27ac and H3K4me1 ChIP-seq data in GM12878. **(D)** An example locus on Chr2:37,300,000-37,300,000 (700Kb) showing (from top to bottom) a chromatin looping domain in 2D contact map of in situ Hi-C data, browser views of CTCF and cohesin ChIA-PET data, ChIP-seq peaks signals demarcating cohesin loading (yellow arrow) and anchoring sites (blue arrow). **(E)** The cohesin ChIA-Drop data in fragment view (top panel) and sorted view (bottom panel) at the same chromatin locus as in (D). The cohesin ChIA-Drop data views were centered at a cohesin loading site measuring single molecule counts depicted for possible one-sided extrusion or two-sided extrusion. **(F)** The CTCF ChIA-Drop data in fragment view (top panel) and sorted view (bottom panel) at the same chromatin locus as in (D). The CTCF ChIA-Drop data views were centered at a cohesin loading site measuring single molecule counts depicted for possible one-sided extrusion or two-sided extrusion.

**Figure S3.**
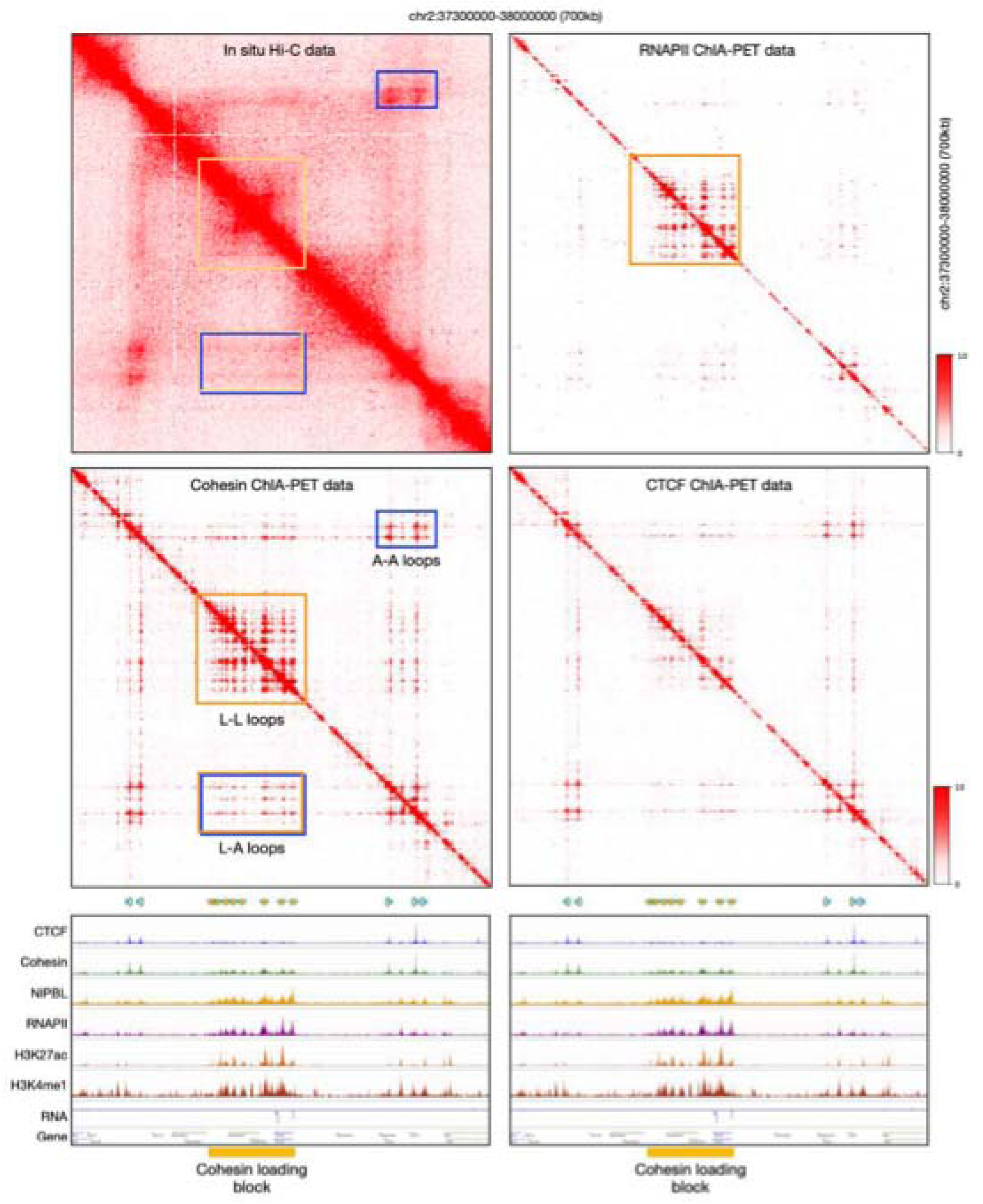
Example of distinct types of chromatin loops (related to Figure 2). Top left: 2D contact map from in situ Hi-C at a representative locus on Chr2:37,300,000–38,000,000 (700 kb). Top right: 2D contact matrix from RNAPII ChIA-PET showing prominent loops between cohesin loading sites (L–L loops, yellow rectangles). Middle left: 2D contact matrix from cohesin ChIA-PET highlighting strong L–L loops, A–A loops (between anchor sites; blue rectangles), and L–A loops (between loading and anchor sites; yellow/blue rectangles). Middle right: 2D contact matrix from CTCF ChIA-PET. Bottom: ChIP–seq tracks for CTCF, cohesin, NIPBL, RNAPII, H3K27ac, and H3K4me1 supporting the identification of the different loop types.

**Figure S4.**
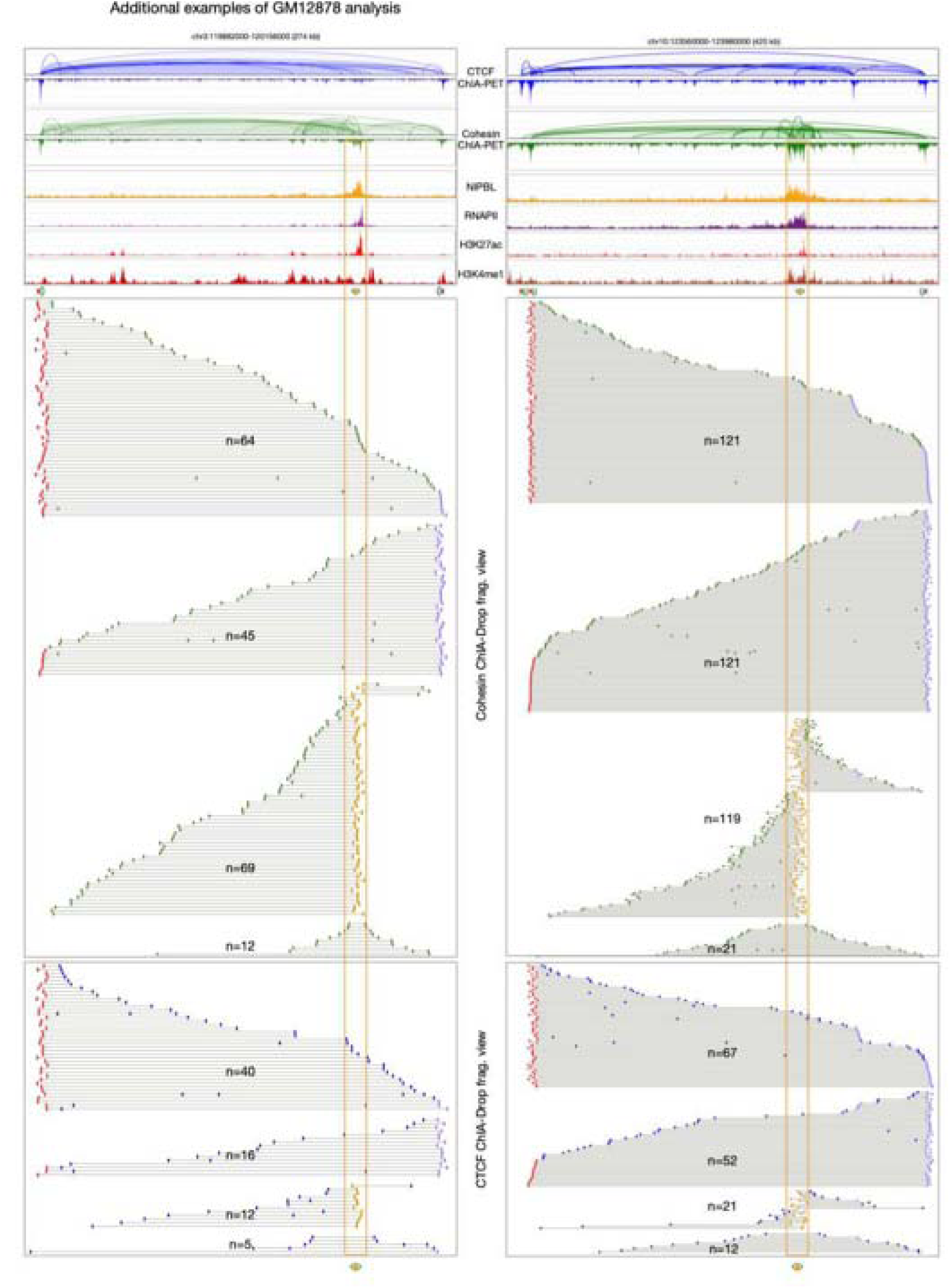
Examples of ongoing and complete chromatin loops (related to Figure 3). Two examples at representative loci: Chr3:119,882,000-120,156,000 (274 Kb) and Chr10:123,560,000-123,980,000 (420 Kb) showing browser view tracks of chromatin loop and peak from CTCF and cohesin ChIA-PET (blue and green) with binding peak tracks from NIPBL, RNAPII, H3K27ac and H3K4me1 ChIP-Seq (top panel); sorted fragment view of chromatin complexes from cohesin ChIA-Drop (middle panel); and CTCF ChIA-Drop (bottom panel). Yellow highlighted rectangular showing the location of cohesin loading site.

**Figure S5.**
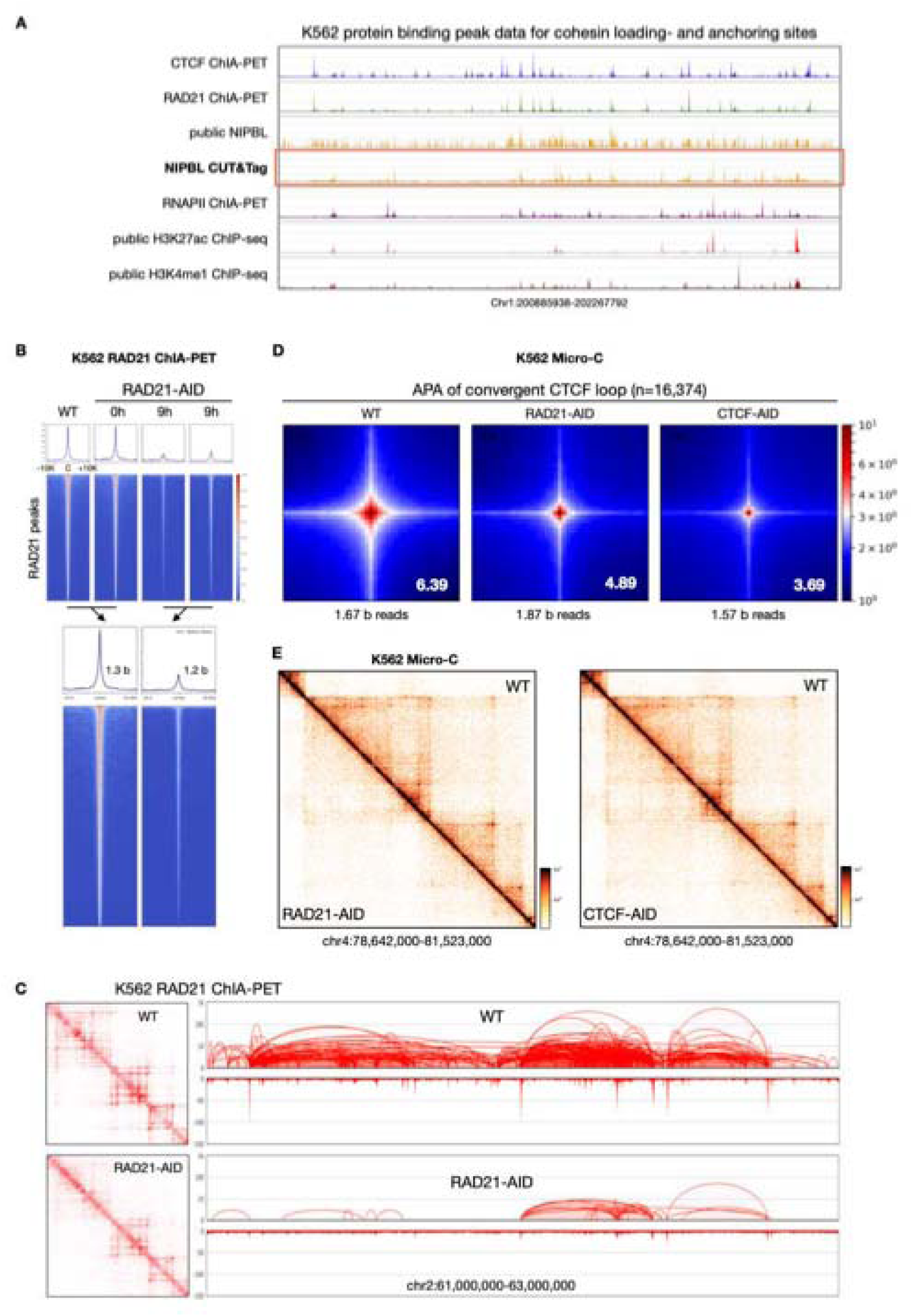
Quality assessment of K562 data for cohesin loading and loop extrusion analysis (related to Figure 4). **(A)** A representative example browser view on Chr1:200,885,938-202,267,792, showing high quality of CTCF, RAD21, NIPBL, RNAPII, H3K27ac and H3K4me1 binding peaks used for the identification of cohesin loading sites and anchor sites. A specifical notion that our NIPBL CUT&Tag data is in high quality while the publicly available NIPBL ChIP-seq data is in very poor quality. **(B)** Aggregation plots of 1D signal at binding sites of RAD21 in wildtype (WT) and no auxin induction control (auxin 0 h) and auxin induction for 9 hours (auxin 9 h). The two control data and the two AID data were combined, respectively, for downstream analysis. **(C)** Example views of 2D contact maps (Left panel) and the browser view (Right panel) fpr RAD21 ChIA-PET data in control (WT) and RAD21-depleted (RAD21-AID) K562 cells at Chr2:61,000,000-63,000,000. **(D)** APA (aggregate peak analysis) of chromatin loops (n=16,374) with convergent (> <) CTCF motifs in control WT (left), RAD21-AID (middle) and CTCF-AID (right) Micro-C data from K562 cells. **(E)** The 2D contact maps of a representative chromatin locus-Chr4:78,642,000-81,523,000 showing the decreased chromatin loops of Micro-C data in RAD21-depleted (RAD21-AID) versus wildtype (WT) K562 cells (left panel) and CTCF-depleted (CTCF-AID) versus wildtype (WT) K562 cells (left panel).

**Fig S6.**
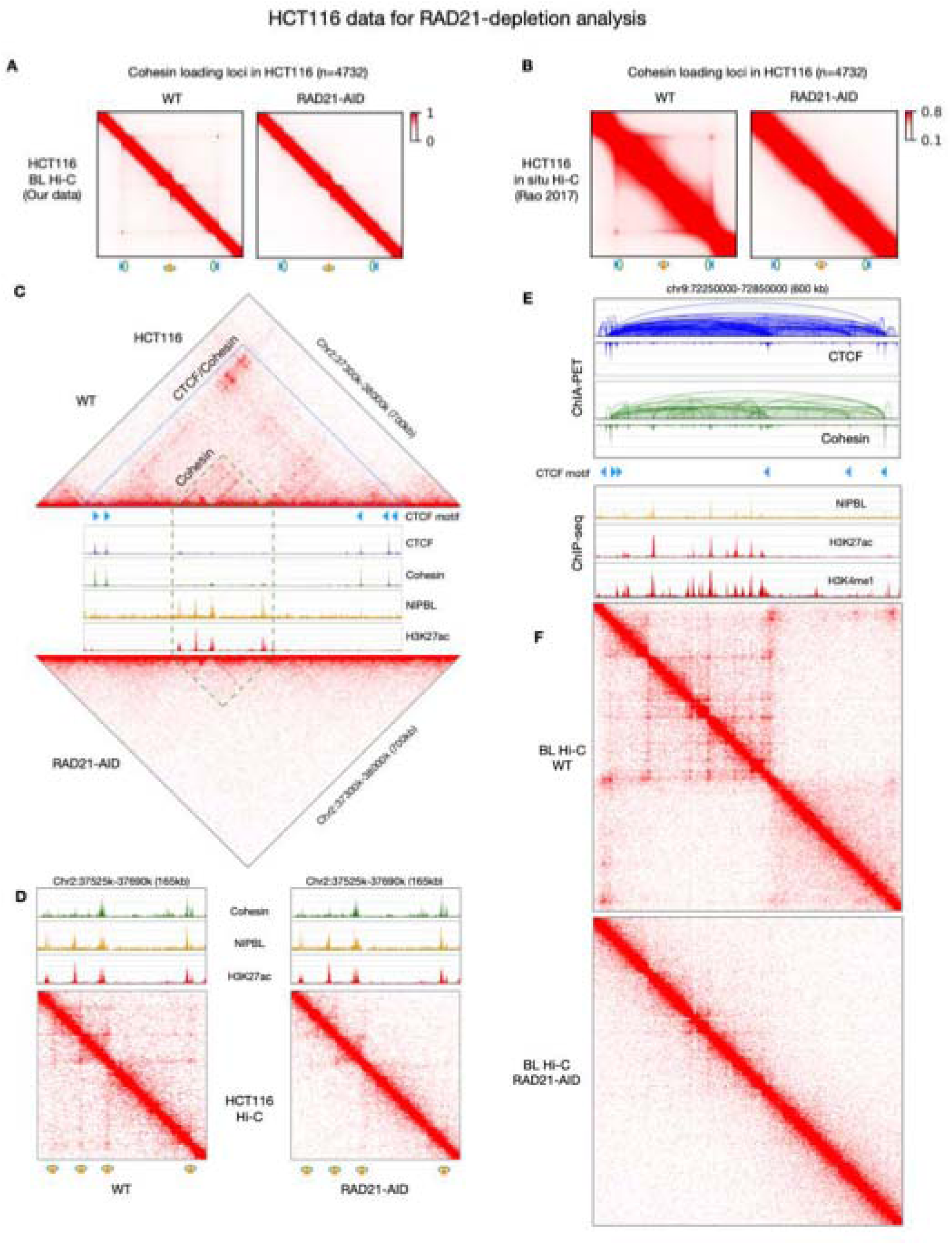
RAD21 depletion in HCT116 cells (related to Figure 4). **(A)** 2D contact maps of aggregating BL Hi-C (non-enriched ChIA-PET or bridge linker Hi-C) data before (WT) and after (RAD21-AID) RAD21-depletion in HCT116 cells centered at the 4,732 cohesin loading sites. **(B)** in situ Hi-C data (Rao, et al, Cell, 2024) before (WT) and after (RAD21-AID) RAD21-depletion in HCT116 cells centered at the 4,732 cohesin loading sites. **(C)** An example of 2D contact triangle maps of a looping domain on chr2 before (WT, top panel) and after (RAD21-AID, bottom panel) depleting in situ Hi-C data, respectively, along with CTCF motifs (blue triangle with orientations); CTCF, cohesin, NIPBL, H3K27ac ChIP-seq tracks (middle panel). The rectangular box with green dashed lines highlighting the region with distinctive cohesin loading sites. **(D)** The zoom-in view of the cohesin loading region showing the cohesin loading-to-loading loops before and after RAD21 depletion. **(E)** An example of chromatin loop domain on Chr9. Top panel, CTCF and cohesin ChIA-PET data defined chromatin loops and peaks; Bottom panel, cohesin loading and anchoring sites demarcated by binding peaks of NIPBL, H3K27ac and H3K4me1 ChIP-seq. **(F)** A 2D contact maps with BL Hi-C data before (WT) and after (RAD21-AID) RAD21 depletion showing mapping signals from individual cohesin sites and loops at the same chromatin loci with figure E.

**Fig S7.**
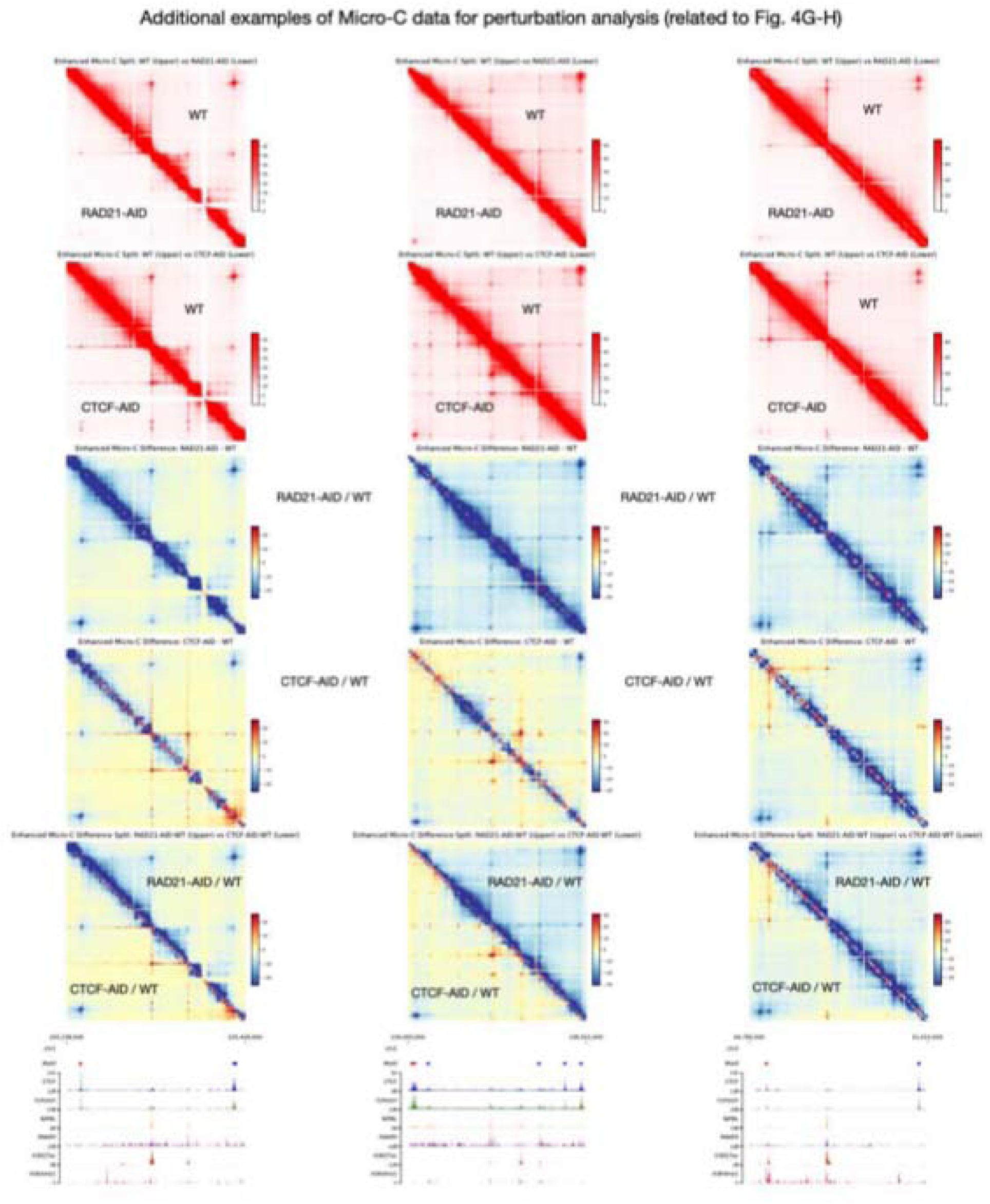
Additional examples for validation of dynamic chromatin looping changes at cohesin loading and anchor sites in RAD21 or CTCF perturbed K562 cells (related to Figure 4G-H). Comparative 2D contact matrix of Micro-C data at three representative chromatin loci: Chr 1: 235,238,000-235,428,000 (left column); Chr 1: 236,093,000-236,501,000 (middle column) and Chr 1: 60,783,000-61,010,000 (right column). Raw 1: RAD21-AID (lower half) vs. WT (upper half). Raw 2: CTCF-AID (lower half) vs. WT (upper half). Raw 3: subtractive 2D contact maps of RAD21-AID / WT. Raw 4: subtractive 2D contact maps of CTCF-AID / WT, Raw 5: CTCF-AID/WT vs RAD21-AID/WT. Raw 6: CTCF, cohesin, NIPBL, RNAPII, H3K27ac and H3K4me1 binding peaks demarcating corresponding cohesin loading and anchoring loci in this domain.

**Figure S8.**
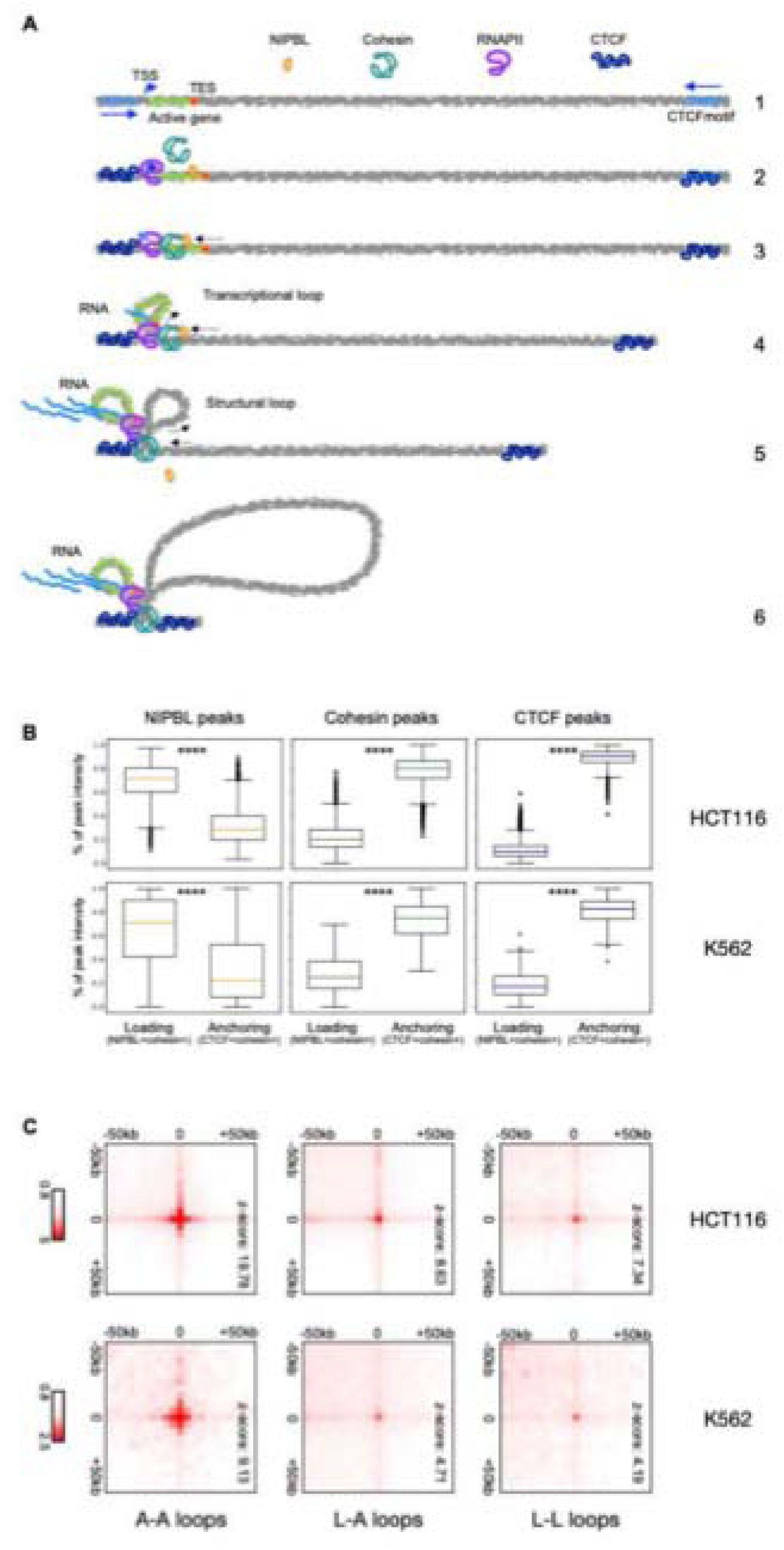
Two phase model of loop extrusion (related to Figure 5). **(A)** Cartoon of the proposed phase II loop extrusion model when NIPBL directly loads proximal to CTCF binding sites. **(B)** Boxplots of binding peak intensity at the cohesin loading and anchoring sites by NIPBL (left), cohesin (middle), and CTCF (right) in HCT116 and K562 cells. ****: *p*-value <. 8e-308. Two-sided Mann-Whitney U test was used for p-values calculation. **(C)** APA analysis for chromatin loop intensity loading-to-loading (L-L), loading-to-anchoring (L-A), and anchoring-to-anchoring (A-A) using in situ Hi-C data in HCT116 and K562 cell lines. Note that the A-A loop intensity is more than two-fold of L-L and L-A loops.

**Figure S9.**
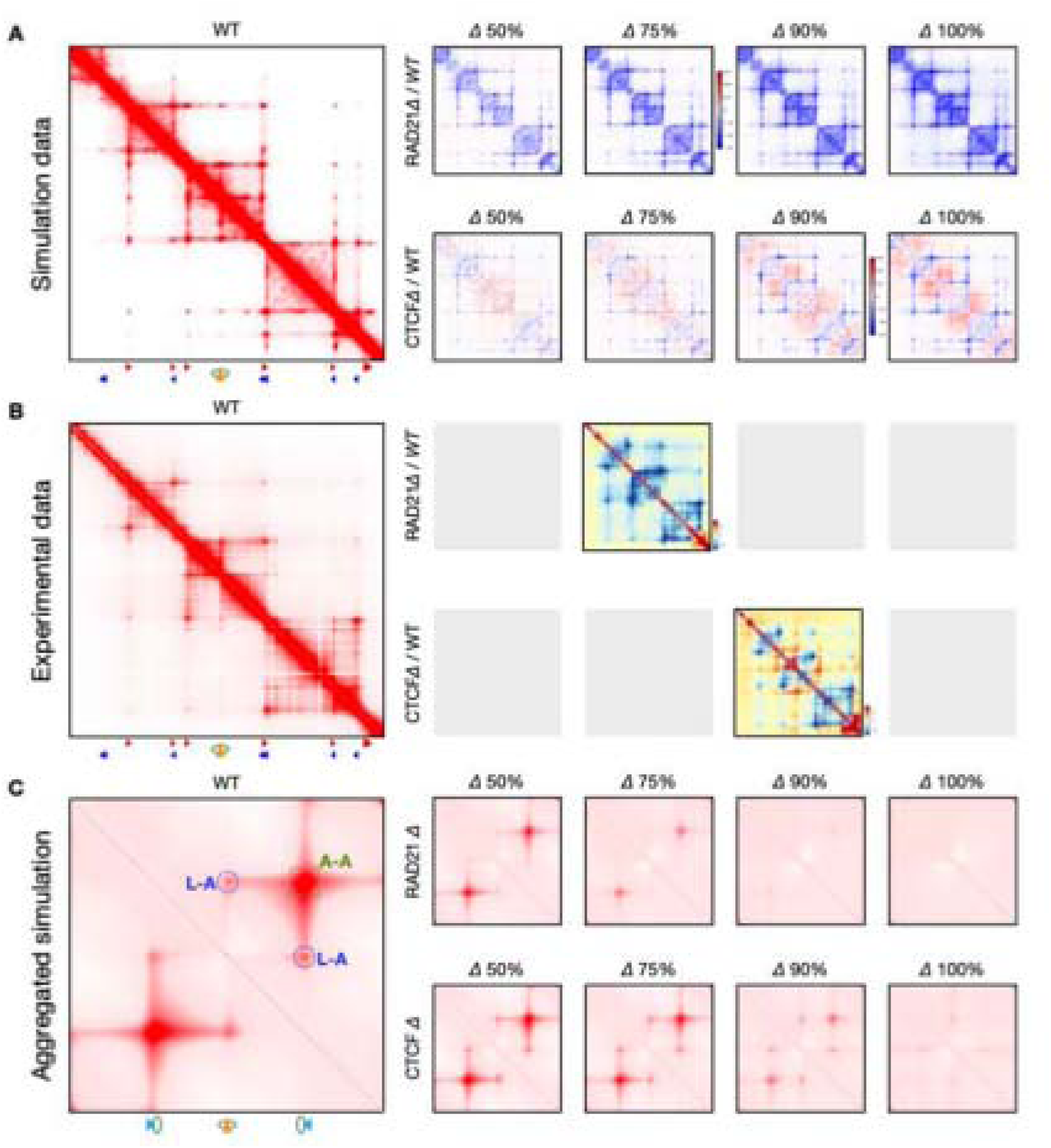
Computational simulations under graded perturbations of CTCF and RAD21 (related to Figure 6). (**A**) Left, Simulated contact map generated by the “One-sided & Two-mode” model for the genomic region “Chr4: 37,302,000-38,102,000” under wild-type (WT) conditions. Right, 4 maps showing changes in contact patterns under increasing levels of RAD21 or CTCF depletion relative to wildtype (WT) condition. (**B**) Experimental Micro-C contact map for the same region in K562 under wildtype (WT) conditions, along with corresponding difference of RAD21 or CTCF depletions that best matched the *in silico* simulated results. Grey squares indicate no match experimental data available. (**C**) Aggregation of distance-normalized simulation maps from 200 genomic regions centered on cohesin loading sites and flanked by convergent CTCF anchor sites, generated using the “One-sided & Two-mode” model. Aggregation plots are shown for wildtype (WT) and graded depletion (Δ) levels of CTCF and RAD21. Position of L-A and A-A loops are indicated by dotted circles.

